# Measuring single neuron visual receptive field sizes by fMRI

**DOI:** 10.1101/301648

**Authors:** Georgios A. Keliris, Qinglin Li, Amalia Papanikolaou, Nikos K. Logothetis, Stelios M. Smirnakis

## Abstract

The non-invasive measurements of neuronal receptive field (RF) properties in-vivo allow a detailed understanding of brain organization as well as its plasticity by longitudinal following of potential changes. Visual RFs measured invasively by electrophysiology in animal models have traditionally provided a great extent of our current knowledge about the visual brain and its disorders. Voxel based estimates of population RF (pRF) by functional magnetic resonance imaging (fMRI) in humans revolutionized the field and have been used extensively in numerous studies. However, current methods cannot estimate single-neuron RF sizes as they reflect large populations of neurons with individual RF scatter. Here, we introduce a new approach to estimate RF size using spatial frequency selectivity to checkerboard patterns. This method allowed us to obtain non-invasive, single-unit, RF estimates in human V1 for the first time. These estimates were significantly smaller compared to prior pRF methods. Further, fMRI and electrophysiological experiments in non-human primates demonstrated an exceptional match validating the approach.

## Introduction

An important contribution of functional magnetic resonance imaging (fMRI) in human neuroscience is the non-invasive in-vivo measurement of the voxel-by-voxel organization of several cortical areas^1^. Recent studies substantially advanced this field of research by using novel neuro-computational methods that can uncover neuronal properties previously only accessible by invasive electrophysiological techniques^2,3^. Estimating neuronal properties in vivo by fMRI is of great significance for understanding the functional organization of the cortex as well as cortical reorganization and plasticity in patients with diseases afflicting the brain^4^.

A prime example of such methods is the estimation of population receptive fields (pRFs) in retinotopically organized visual areas^5–7^. However, pRFs are only estimates of aggregate voxelbased averages of ten to hundreds of thousands of neurons within fMRI voxels and are a function of: a) the receptive field properties of single units belonging to a voxel, b) the scatter in the location of receptive field centers across units, and c) the interactions between nearby connected units. Here, we present a novel approach to estimate, for the first time, the average single-neuron receptive field sizes in human primary visual cortex. To this end, we exploit the spatial-frequency dependent fMRI responses of visual RFs modeled as Gabor functions. Furthermore, we validate non-invasive RF size estimates obtained using the same fMRI method in non-human primates by comparing them directly with RF sizes obtained via intracranial electrophysiological recordings.

The retinotopic organization of visual cortex has been extensively studied in primates. Very early on, studies indicated that a key property of early striate and extrastriate areas is the expanded cortical representation of the central visual field often called magnification factor^8–10^. More recently, this led to the development of analytical formulations to describe the projection from the visual field (retina) to the cortical space^11–13^. Typically, the spatial sampling of fMRI happens uniformly in cortical space with voxel sizes of the order of 1^3^−3^3^ mm^3^. This, in combination to the non-linear mapping of visual to cortical space, has direct implications for estimating population receptive field sizes. To demonstrate that, we used the inverse of the simple k·log(z+a) model described by Schwartz (1977) to transform the coordinates of square pixels from cortical space to visual space (Fig. 1 A-B). Figure 1B shows the expected RF sizes of neurons whose RF centers lie along the edges of the transformed fMRI voxels (red circles) given no random scatter^14^. Dashed circles represent the envelopes of each voxel’s expected pRF size. Note that expected pRF size, strongly depends on the non-linear transformation from cortical to visual space, clearly overestimating individual neuron RF size.

**Figure 1.**
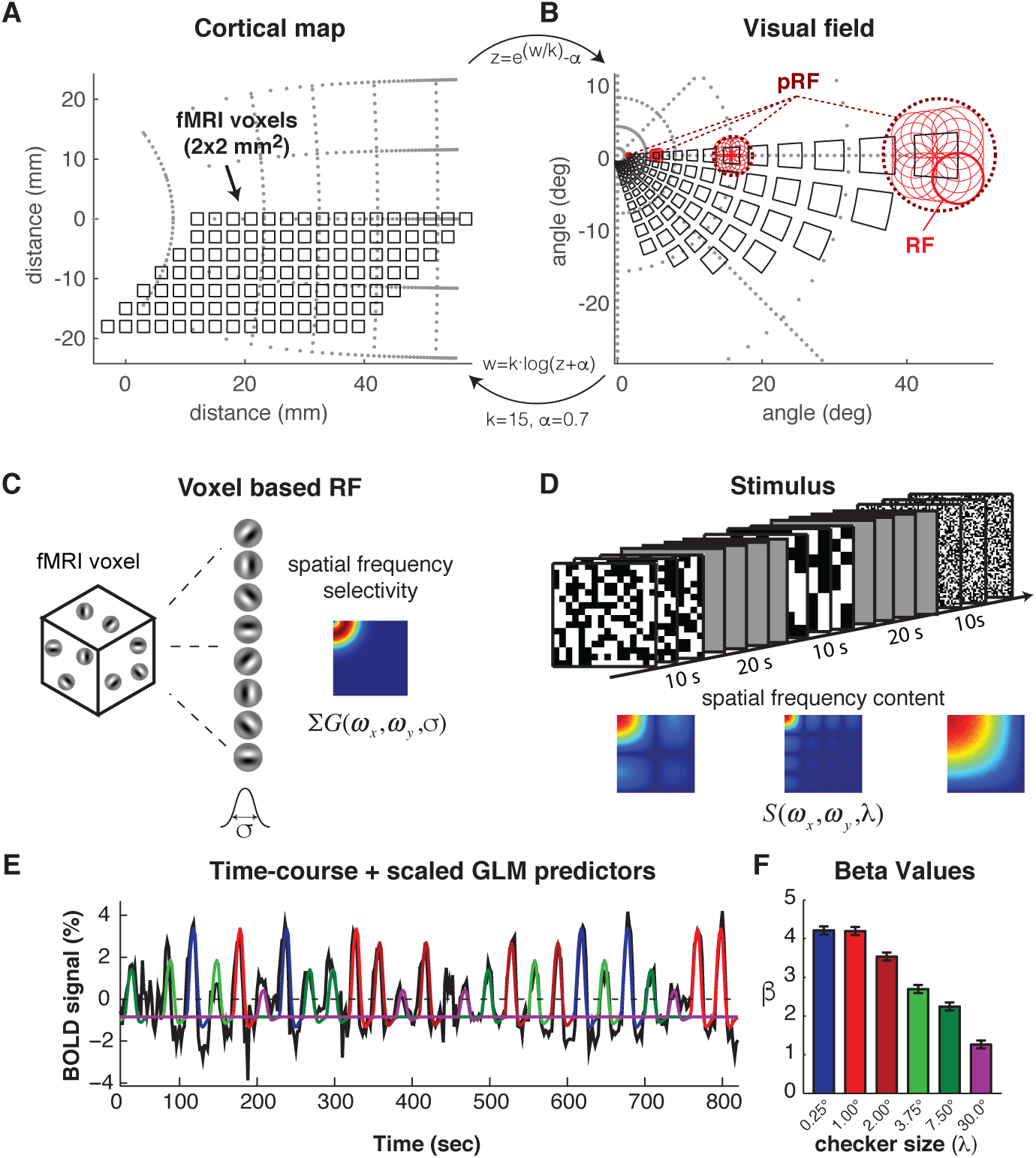
Visual field back-projection of fMRI voxels, RF modeling approach, and example of GLM fit of stimulus predictors. (A-B) Transformation of the fMRI-voxels (squares) from the cortical space (A) to the visual field (B) by using the inverse-Schwartz model^11^. In (B) we plot estimated RF sizes along voxel boarders at different eccentricities (red circles). Population RF size (pRF) depends not only on RF size but also on the non-linear transformation from cortical to visual field. (C) We modeled suRFs as 2D-Gabor functions assuming that voxels contain an approximately homogenous representation of all orientations. Thus, voxel spatial frequency selectivity becomes independent of orientation and can be estimated according to the *σ* of the Gabor envelope (see Fig S1). (D) Full-field checkerboard patterns were presented in blocks of 10 seconds ON, 20 seconds OFF. Blocks were pseudo-randomly interleaved across six different spatial frequency conditions (checker sizes; see Fig S2). (E) Raw BOLD-signal timecourse from a sample region of interest in area V1 of one subject (black) overlaid by the GLM-fit of the six predictors (checker sizes) in different colors. (F) The GLM (β-weights corresponding to each checker size predictor.

Receptive fields of neurons in primary visual cortex are thought to act as spatial frequency filters^15^ and can be approximated by 2D-Gabor functions^16^ (see Fig. S1). The spatial frequency selectivity of these filters can be analytically estimated depending on Gabor function parameters including its Gaussian envelope’s standard deviation, which is proportional to the RF size (Fig. 1 C). We hypothesized that fMRI responses to stimuli with different spatial frequency contents could be used to estimate the underlying neuronal Gabor function standard deviations based on their spatial frequency selectivity and thus independent of spatial position and scatter. To this end, we chose to use binary white noise checkerboard patterns as stimuli, modulating their spatial frequency content by changing the checker size across conditions (Fig. 1 D). We note that our Gabor based modeling approach would also be valid, if we used other stimuli with manipulated or selected spatial frequency content like sinewave gratings or filtered natural images. Here we used checkerboard patterns since their frequency content could be estimated analytically allowing easier computational modeling (Fig. S2).

## Results

Responses in human primary visual cortex were nicely modulated by our stimuli (Fig. 2). On average, V1 responded more strongly to blocks with smaller checkers (Fig. 2 B-C). More specifically, as expected from our hypothesis, voxels selected from anatomical and functionally defined areas close to the fovea were biased to the smallest checker sizes while gradually more peripheral voxels showed stronger responses to larger sizes (Fig. 2 D-G).

**Figure 2.**
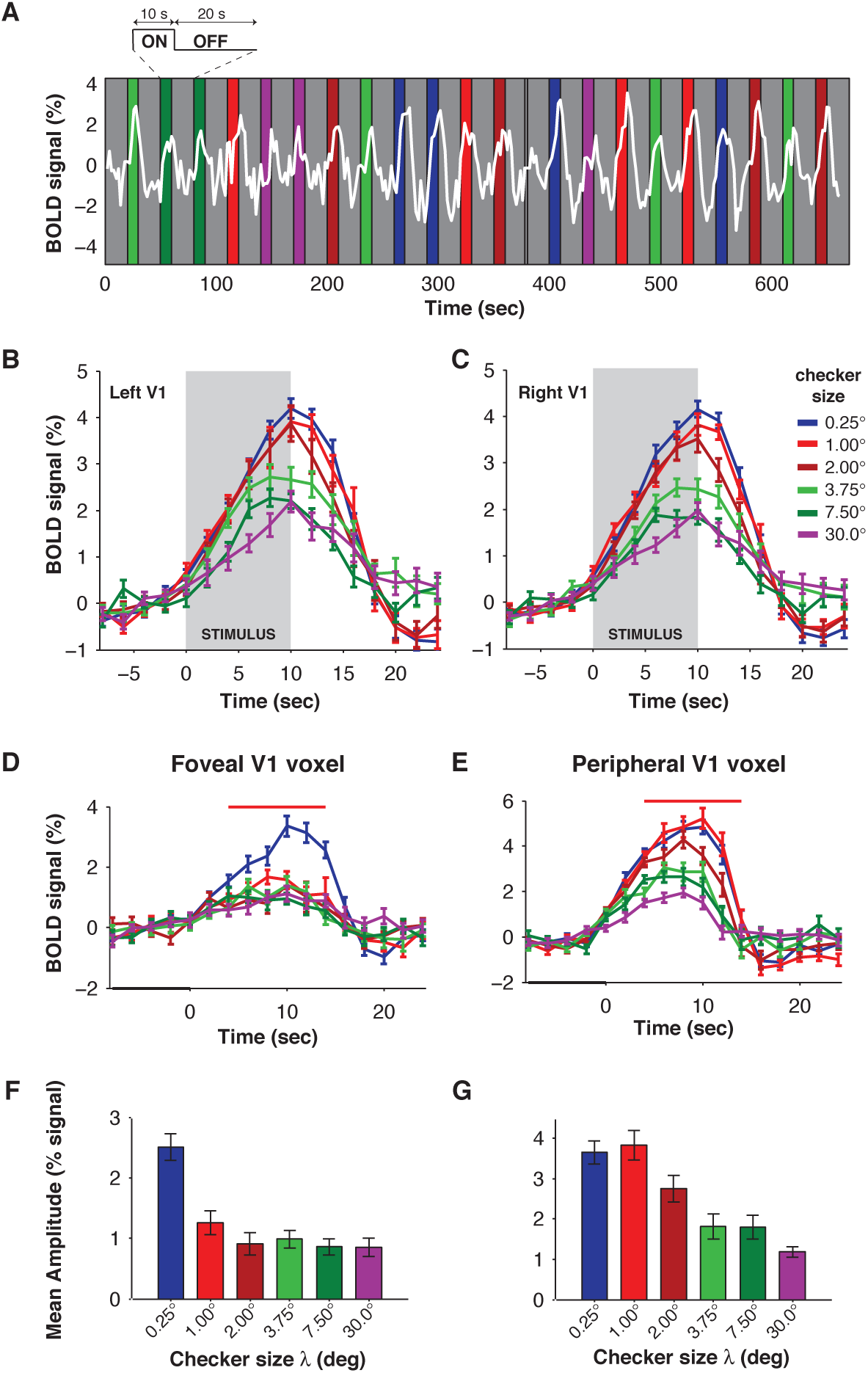
Example of paradigm and V1 responses. (A) An example of the fMRI response in V1 of a subject is shown on top of a cartoon of the stimulus presentation with different conditions in colored blocks (10 seconds) and the inter-stimulus interval (20 seconds) in gray. (B-C) The event related time courses for each condition are shown for left (B) and right V1 (C). (D-E) Similarly, the event-related time courses of selected single voxels in foveal (D) and peripheral (E) V1 are shown. (F-G) represent the mean amplitude of the BOLD signals during a window around the peak (red horizontal bar in D and E).

To estimate the average single-unit RF (suRF) size in a voxel we used model-types of the form:

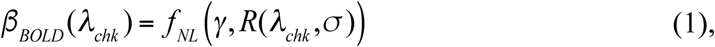

where *β_BOLD_*(*λ_chk_*) is the GLM beta parameter that reflects the response to the stimulus block with checker size *λ_chk_*,*f_NL_* is a static non-linearity, *γ*a gain parameter and *R* the estimated neural response as a function of stimulus condition *λ_chk_* and the Gabor filter parameters. *R* is given by:

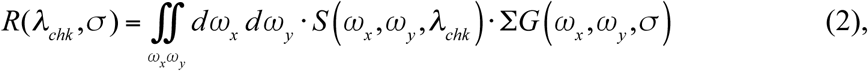

where *S*(*ω_x_*,*ω_y_*,*λ_chk_*) represents the spatial frequency content of the stimulus (see Fig. S2), and Σ*G*(*ω_x_*,*ω_y_*, *σ*) the spatial frequency selectivity of a voxel (see Fig. S1). We considered RF-size to be equal to 2*σ*, as this is close to the classical way of mapping RFs by detecting the first response to moving stimuli across the RF-edges.

Several models of the above type were tested and in particular ones using a log-square-root compressive (LSC) non-linearity or a power-low compressive non-linearity (PLC); some models had divisive (dPLC) or subtractive (sPLC) surround components (Table 1). For a qualitative overview of model behaviour across a range of values in parameter space see Fig. S3–S6. All models demonstrated exceptionally good fits as reflected in very high median coefficients of determination (CoD) across all subjects (Table 2). To select the best model we also calculated Akaike’s information criterion (AIC) and from there derived the relative likelihoods of each model per voxel^18^. Based on the mean AIC values (Table 2), and the distributions of relative likelihoods (Fig. S7) we chose the PLC model, which simulates only the RF-center, for further reporting our results. Models with divisive or subtractive surround components on average only marginally improved CoD or did not show improvements and had slightly worse AIC scores due to the higher number of parameters (Table 2). It should be noted, however, that a small number of voxels demonstrated clear suppressive effects for intermediate checker-size conditions with the center only model not able to account for this suppression. In those voxels, models that included the surround and in particular the sPLC were clearly outperforming the center only model (Fig. S8).

**Table 1.** Models used for fitting the Gabor receptive field parameters by using fMRI responses. The parameters *β* are GLM estimates of the fMRI response amplitude for each stimulus condition, *γ_c_* and *γ_s_* are gain parameters for center and surround (when applicable) respectively, and similarly *R*(*σ_c_*), *R*(*σ_s_*) are the estimated neural responses of gabor-like neurons with Gaussian envelope with standard deviation *σ* as given by equation (2) in the main manuscript. The exponent parameter *n* acts as a compressive non-linearity and was estimated to be around 0.325 by Kay et al. 2013^17^. For dPLC and sPLC we have set it to this estimate to avoid over-fitting due to a high number of parameters.

| Model ID | Equation | Name |
| --- | --- | --- |
| LSC | $\beta = \gamma_c \cdot \log\left(1 + \sqrt{R(\sigma_c)}\right)$ | Log-Square root-Compressive |
| PLC | $\beta = (\gamma_c \cdot R(\sigma_c))^n$ | Power-Law-Compressive |
| dPLC | $\beta = \gamma \cdot \left( \frac{\gamma_c \cdot R(\sigma_c)}{1 + \gamma_s \cdot R(\sigma_s)} \right)^{0.325}$ | Divisive normalization PLC |
| sPLC | $\beta = (\gamma_c \cdot R(\sigma_c))^{0.325} - (\gamma_s \cdot R(\sigma_s))^{0.325}$ | Subtractive normalization PLC |

**Table 2.** Evaluation of model performance in human subjects. Performance was calculated based on the median coefficients of determination (CoD) as well as Akaike’s information criterion (AIC)^18^. Based on these values and relative likelihoods of the models (see Fig. S8) we selected the PLC model (in BOLD font) for presentation of the results in the body of the manuscript. Abbreviations: LSC: Log-Square root-Compressive, PLC: Power-Law-Compressive, dPLC: devisive normalization PLC, sPLC: subtractive normalization PLC.

| Model ID | # params |  | H1 | H2 | H3 | H4 |
| --- | --- | --- | --- | --- | --- | --- |
| LSC | 2 | CoD<br>AIC | 98.4 %<br>-3.6754 | 97.8 %<br>-3.4150 | 97.3 %<br>-3.0358 | 97.7 %<br>-3.3004 |
| **PLC** | **3** | **CoD**<br>**AIC** | **98.9 %**<br>**-3.9078** | **98.8 %**<br>**-3.7378** | **98.5 %**<br>**-3.4641** | **99.1 %**<br>**-3.9815** |
| dPLC | 5 | CoD<br>AIC | 99.1 %<br>-3.4779 | 99.1 %<br>-3.3309 | 98.6 %<br>-2.8225 | 99.1 %<br>-3.3747 |
| sPLC | 4 | CoD<br>AIC | 98.5 %<br>-3.1736 | 98.0 %<br>-2.8642 | 98.0 %<br>-2.8409 | 98.1 %<br>-2.9591 |

Examples of the PLC-suRF model fit for each subject H1-H4 are presented in Fig. 3 A-D respectively. On the top row, voxels with eccentricity closer to the fovea were selected based on the classical pRF model fit (Methods) and gradually higher eccentricities in the middle and bottom rows. As expected from model simulations (see Fig. S3 C), the shape of the model fit (solid lines) is predicting small RF sizes for foveal voxels and gradually larger for the more peripheral ones. To better demonstrate the relationship between estimated voxel-based RF sizes and eccentricity we performed linear regression for each subject (Fig. 3 E-H; **black lines**). For comparison, linear regression was also performed for the pRF model across the same subjects (Fig. 3 E-H; **gray lines**). The results demonstrated a significant linear relationship of suRF as well as pRF size with eccentricity for all individual subjects and across the population (**see** Table S1) To test if the suRF estimated RF sizes were smaller than the pRF, as we hypothesized (see Fig. 1), we performed analysis of covariance (ancova) and second level comparisons (Tukey-Kramer) of the intercepts (suRF: 0.69 ± 0.07 [95%CI], pRF: 0.94 ± 0.19) and slopes (suRF: 0.05 ± 0.01, pRF: 0.33 ± 0.02) of the two models across the population (Hl-4). We observed a significant difference of the intercepts (*P*=0.01) as well as the slopes (*P*= 1.06×10^−10^), being smaller for the suRF model as expected.

**Figure 3.**
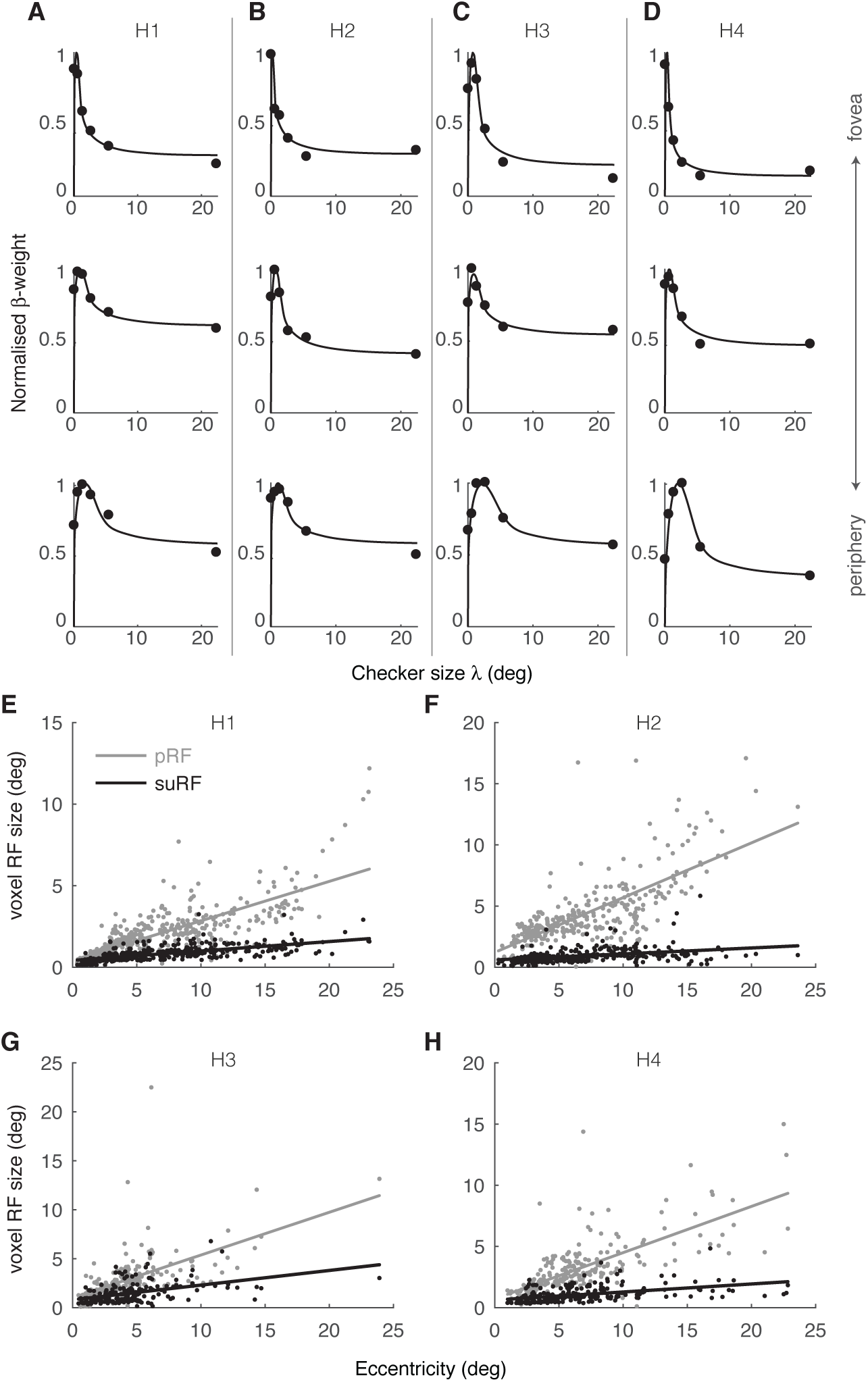
Comparison of suRF with pRF. (A-D) Examples of suRF voxel fits for subjects H1-H4. Each colmnn presents example voxel responses ((β-weights per condition; dots) from gradually increasing eccentricity (top to bottom). Solid lines represent PLC-suRF model fits. (E-H) suRF size (black) and pRF size (gray) as function of eccentricity for subjects H1-H4. Lines respectively represent linear regression fits.

To better understand the relationship between our proposed suRF model and electrophysiological measurements of RF sizes, we performed additional experiments in rhesus macaques. During anesthetized fMRI experiments monkeys were presented with identical stimuli as human subjects. Further, pRF as well as suRF models were estimated with the same methodology as used for the humans (Methods). As shown in the example voxels in Fig. 4 A-D, the responses and suRF-model fits for monkeys were very similar to humans with lower eccentricity voxels demonstrating smaller RF sizes in comparison to voxels located at more peripheral locations. Linear regression analyses of the eccentricity versus suRF and pRF sizes demonstrated identical results as the in humans (Fig. 4 E-F; Fig. S10) with all subjects showing significant linear relationships for both models (see Table S2). Furthermore, we performed ancova followed by the comparison (Tukey-Kramer test) of intercepts (suRF: 0.42 ± 0.05, pRF: 1.39 ± 0.15) and slopes (suRF: 0.06 ± 0.01, pRF: 0.19 ± 0.02). As in humans, we observed significant differences of both intercepts (*P*=9.56×10^−10^) and slopes (*P*=9,56×10^−10^), Importantly, the estimated suRF sizes approximated previously reported electrophysiological measurements of single unit RFs^10,19,20^.

To more directly investigate how suRF-fMRI estimates compare with single neuron electrophysiological RF sizes, we then performed electrophysiological RF measurements in two other monkeys (M3-M4). To this end, we have used the method of reverse correlation that has been extensively used in previous RF-mapping experiments in visual cortex^21–25^. In addition, we also used a moving bar method that closely resembles the pRF mapping we used in fMRI (Methods). Examples of the recorded RFs are presented in Fig. 4H and Fig. S11-**S16**. Since the electrophysiology data from both monkeys and methods were consistent we have collapsed them and performed linear regression for RF size versus eccentricity like we did for the fMRI measurements, which are presented in the same figure (Fig. 4 G). Comparison of the suRF intercept and slope with electrophysiology (intercept=0.16 ± 0.05, slope=0.08 ± 0.02) showed no significant difference (intercepts: *P*=0.47, slopes: *P*=0.98; See Fig. 4 G and **Table S4**).

**Figure 4.**
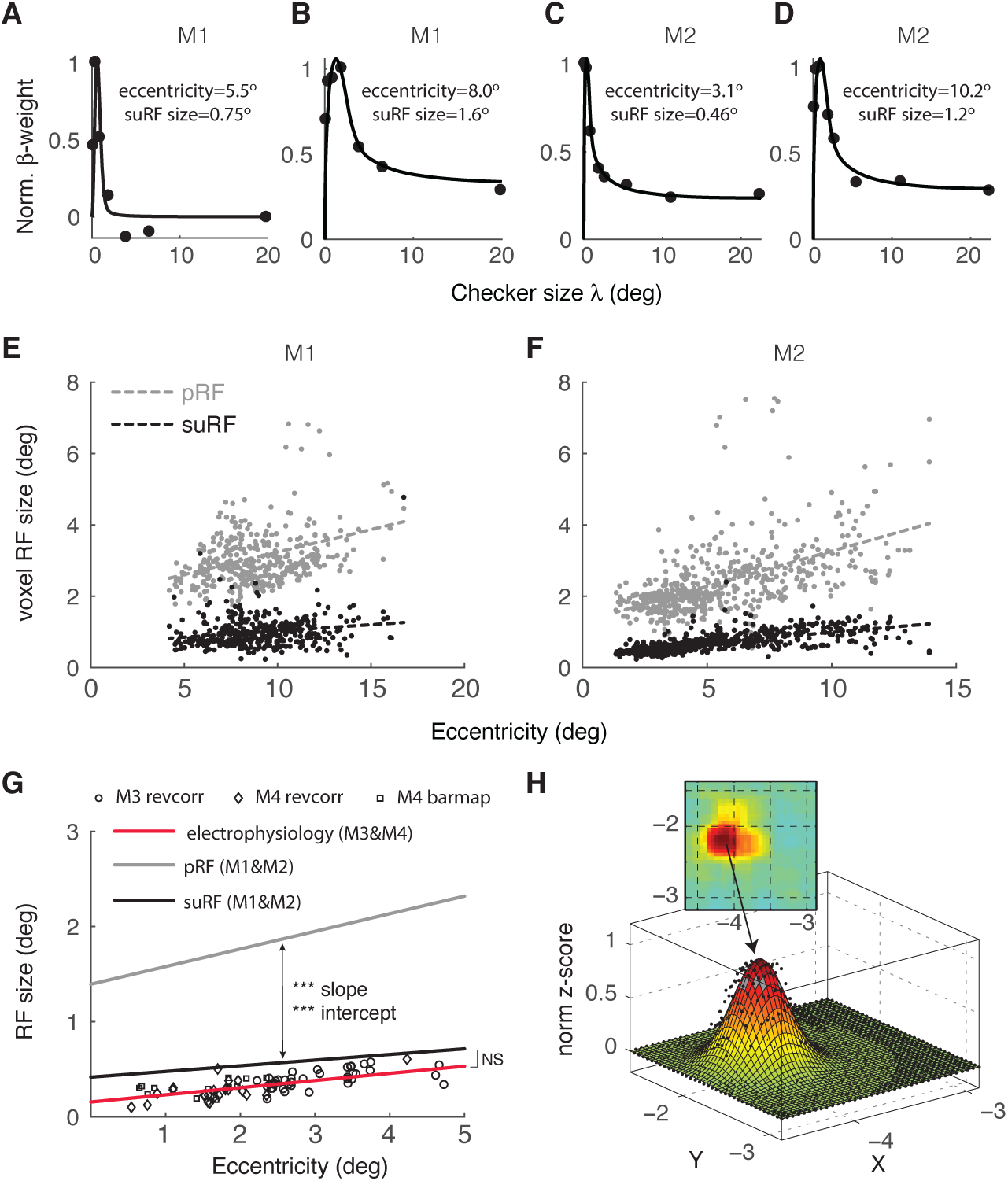
Validating suRF with electrophysiology in Rhesus Macaques. (A-D) Example voxel responses (β-weights) and respective suRF fits (solid lines) for monkeys M1-M2. (E-F) pRF (gray) and suRF (black) size as a function of eccentricity for monkeys M1-M2 (see also Fig. S10). Each dot represents a voxel. Dashed lines: linear regression fits (see Table S4 for statistics). (G) pRF and suRF fits across both monkeys M1&M2 compared to electrophysiology obtained from two other animals (M3&M4; red). For parameter estimates and statistics see Table S4. (H) Example of electrophysiology RF estimate by reverse correlation (see also Fig. S11**-S16**). Abbreviations: ^***^p<10^−9^, NS p>0.05

To absolutely settle the correspondence between the suRF model and electrophysiology, we performed fMRI pRF and suRF experiments in monkey M3 that had MRI compatible implants (Fig. 5). To be able to coregister the physiology and MRI estimates we have inserted an MRI compatible guide (Fig. 5 B) on top of the grid in the chamber and filled it with an MRI contrast agent (MION). In this way, a reference frame was reconstructed and we could use it to estimate the voxel corresponding to our recording electrode. In Fig. 5 C-E, we show estimates of the RF size based on all three methods (pRF, suRF, electrophysiology). As one can appreciate there is a direct close correspondence between the electrophysiology and suRF estimates that are both much smaller in comparison to the pRF (Fig. 5 F).

**Figure 5.**
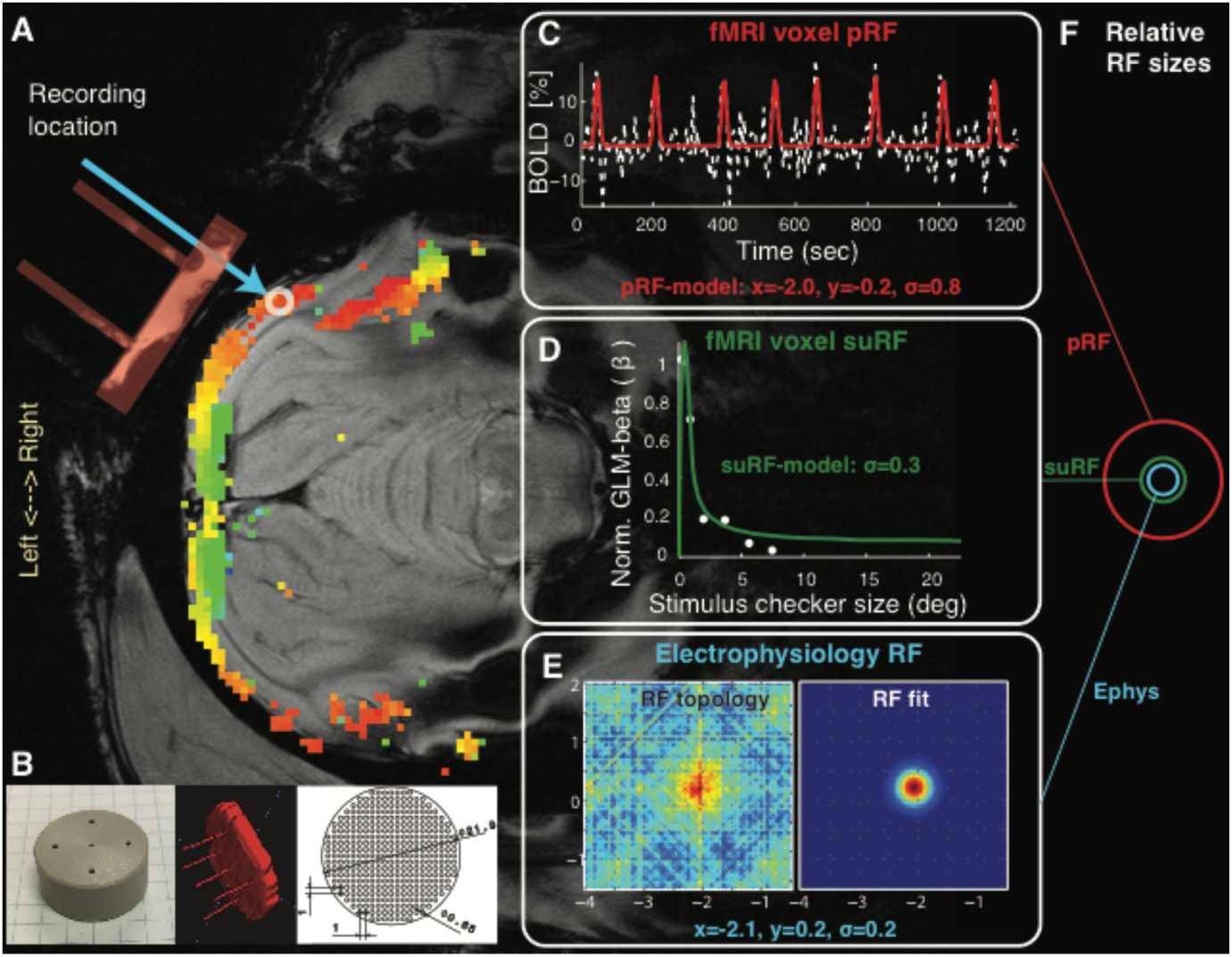
Comparison of suRF with pRF and electrophysiology in the same animal. (A) Anatomical slice from the recording location with overlaid pRF eccentricity map. To localize the recording location, we used an MRI compatible insert (B; left) in the implanted recording chamber on top of the already present recording grid (B; right). The chamber including the grid and holes of insert were filled with MRI contrast agent so we could reconstruct their position (B; middle) and localize the position of the electrode recordings that happened in separate sessions outside the scanner (light blue arrow) to avoid artifact. (C) The pRF model fit and parameters are plotted on the fMRI signal timecourse from a voxel under the electrode tip (white circle). (D) suRF model fit for the same voxel. (E) Electrophysiology RF from the same location using the moving bar method (see Methods). (F) Relative RF size for all three methods. Note suRF closely approximates the “ground truth” of the electrophysiology RF size.

## Discussion

To date, only a couple of studies attempted to report measurements of receptive field sizes in human visual cortex either with invasive intractranial electrophysiology (potentially 1-2 cells)^26^ or surface electrocorticography in patients^27^. Here, we reported and validated by electrophysiology in macaques, estimates of single unit RF sizes in human primary visual cortex by using *in*-*vivo* fMRI. To the best of our knowledge, our study provides the first comprehensive measurements over the whole primary visual cortex using a simple technique that can be easily used in the wide population.

The non-invasive fMRI suRF method provides exceptionally good estimates of V1 RF sizes that closely match invasive electrophysiological measurements. Further, the results in humans were closely matched with those in rhesus macaques in which we validated the methodology. Thus, the use of fMRI in both species provides a necessary bridge for human fMRI research to the gold standard and allows further non-invasive studies of human cortical reorganization in the case of injury or disease.

## Methods

### Human Experiments

#### Subjects

Four healthy human subjects (H1-H4, 24-44, 2 females) with normal or corrected-to-normal visual acuity participated in the fMRI experiments. Before each session subjects provided written informed consent. The local ethics committee of the University Hospital Tübingen approved the study.

#### MRI data acquisition and preprocessing

MRI experiments were performed at the Max Planck Institute for Biological Cybernetics, Tübingen, Germany. Functional and anatomical images were acquired in a 3.0 Tesla Tim Trio Scanner (Siemens Ltd., Erlangen, Germany) using a 12-channel coil. At least two T1-weighted anatomical volumes were acquired for each subject with a 3D magnetization prepared rapid acquisition gradient echo (T1 MPRAGE scan) and averaged following alignment to increase signal to noise ratio (matrix size=256×256, voxel size=1×1×1 mm^3^, 176 partitions, flip angle=9°, TR=1900 ms, TE=2.26 ms, TI=900 ms). Blood oxygen level dependent (BOLD) image volumes were acquired using gradient echo sequences of 28 contiguous 3 mm-thick slices covering the entire brain (TR=2000 ms, TE=40 ms, matrix size=64×64, voxel size=3×3×3 mm^3^, flip angle=90°).

At least 5 functional scans were acquired for each subject, consisting of 195 image volumes, the first 3 of which were discarded. The functional images were corrected for motion between and within scans^28^ and were aligned to the high-resolution anatomical volume using a mutual information method^29^. The high-resolution anatomical data were used to segment the white-gray matter boundary in itkGray software, and 3D cortical surface and flat mesh models were created and realigned with the functional data. The functional time-series were spatially resampled in the volume space using nearest neighbor interpolation. This preserves the original signals but up-samples them in space leading to some voxels with the same time-courses. Analysis was accelerated by analyzing a single voxel corresponding to the original fMRI resolution and assigning the results to the corresponding anatomical voxels. All subsequent analysis was performed in the segmented volume space restricted in the gray-matter voxels. The above preprocessing steps were performed in MATLAB using the mrVista software package that can be found at https://github.com/vistalab/vistasoft.

#### Population receptive field (pRF) mapping

For retinotopic mapping we used the population receptive field (pRF) method^5^. Shortly, the pRF model estimates the region of the visual field that effectively elicits a response in a small region of visual cortex (voxel). The implementation of the pRF model is a circularly symmetric Gaussian receptive field in visual space, whose center and radius are estimated by fitting actual BOLD signal responses to estimated responses elicited by convolving the model with the moving bar stimuli. We retained only those voxels in the visual area, for which the pRF model explained more than 15% of the variance. This threshold was set after measuring the mean explained variance in a non-visually responsive area and setting the threshold at six standard deviations above the mean. This method derived reliable and reproducible retinotopic and pRF size maps.

### Stimuli

#### Stimulus presentation

Subjects were presented with visual stimuli in the scanner by using MRI compatible digital goggles (VisuaStim, Resonance Technology Company, Inc, Northridge, CA, USA; 30° horizontal and 22.5° vertical field of view, 800×600 resolution, minimum luminance = 0.3 cd/m^2^ and maximum luminance = 12.2 cd/m^2^).

#### Population RF (pRF) mapping

Moving square-checkerboard bars (100% contrast) were presented within a circular aperture with a radius of 11.25° (full vertical extent of screen) around the fixation point. The bar width was 1.875° and travelled sequentially in 8 different directions, moving by a step half of its size (0.9375°) every image volume acquisition (TR=2 seconds). Stimuli were generated using Psychtoolbox-3^30^ and an open toolbox (VISTADISP) in MATLAB (The Mathworks, Inc.). The subjects’ task was to fixate a small dot in the center of the screen (radius: 0.0375°; 2 pixels) and respond to the color change by pressing a button. The color was changing randomly with average frequency of one every 6.25 seconds. An infrared eye tracker was used to record eye movements inside the scanner (iView XTM, SensoMotoric Instruments GmbH).

#### Single unit RF (suRF) mapping

Binary full-field checkerboards (black/white; 100% contrast) with a frame rate of 30 Hz were presented in a block design (10 seconds ON, 20 seconds OFF). For each block the size (side) of the checkers was chosen to be one of six possible conditions [0.25°, 1°, 2°, 3.75°, 7.5°, 22.5°].

### Monkey Experiments

#### Subjects

Four healthy adult rhesus monkeys (*macaca mulatta*; M1-M4, 5-11 kg, 1 female) were used in MRI (N=3, M1-3) and electrophysiology experiments (N=2, M3-4). The experimental and surgical procedures were performed with great care and in full compliance with the German Law for the Protection of Animals, the European Community guidelines for the care and use of laboratory animals (EUVS 86/609/EEC), and the recommendations of the Weatherall report for the use of non-human primates in research. The regional authorities approved our experimental protocols and the institutional representatives for animal protection supervised all procedures. Animals were kept in large cages located adjacent to the training and experimental facilities. Space in these cages allows swinging and jumping, and enrichment equipment such as toys was changed frequently. Group housing was maintained to increase the quality of life by rich visual, olfactory, auditory and social interaction and stimulation for play. Balanced nutrition and regular veterinary care and monitoring, were provided. Recording chamber implantations were performed in two animals while the animals were under general anesthesia and aseptic conditions. To alleviate post-surgical pain, we administered analgesics for a week after the surgery (see surgical procedures below). MRI experiments were also performed under anesthesia (see anesthesia below). Animals were not sacrificed after the experiments.

#### Anesthesia

Details on the anesthesia protocol have been given previously^31^. Briefly, the animals were premedicated with glycopyrolate (0.01 mg/kg, intramuscular (IM)) and ketamine (15 mg/kg, (IM)) and then deep anesthesia was induced by fentanyl (3 μg/kg), thiopental (5 mg/kg), and succinyl chloride (3 mg/kg). Anesthesia was maintained with remifentanyl (0.5-2 μg/kg/min) under paralysis with mivacurium chloride (3-6 mg/kg/h) to ensure the suppression of eye movements. Heart rate and blood oxygen saturation were monitored continuously with a pulseoxymeter. Body temperature was kept constant at 37-38 °C.

#### Surgical procedures

Recording chambers were positioned over the operculum in area V1 according to stereotaxic coordinates. This was aided by high-resolution magnetic resonance anatomical imaging. The anatomical scan and recording chamber implantation were done while the animals where under general anesthesia. Details of the procedure can be found in our previous work^31^.

#### MRI data acquisition and preprocessing

FMRI experiments were performed in a 4.7T or 7.0T vertical scanner (Bruker Biospec, Bruker Biospin GmbH, Ettlingen, Germany). Multislice fMRI was performed using 8-segmented gradient-recalled echo-planar imaging (EPI). Volumes of 17 slices of 0.756×0.756×2 mm^3^ were collected, each with a field of view (FOV) of 96 × 96 mm on a 128 × 128 matrix and 2 mm slice thickness, flip angle (FA) 40° for 4.7T and 47.6° for 7T, echo time (TE) 20 ms and a repetition time (TR) of 750 ms per segment resulting in a volume acquisition time of 6 seconds. For anatomical measurements, we used a FLASH sequence with the same FOV 96 × 96 mm^2^, matrix 256×256, slice thickness 2 mm, flip angle=70°, TE=10 ms and TR=2000 ms. A highresolution 3D-MDEFT anatomical scan with an isotropic resolution of 0.5 mm^3^ was acquired for co-registration with the FLASH and EPI images. For more details on the fMRI acquisition methods see previously published papers^31,32^. Then, fMRI data were reconstructed and imported into a MATLAB based toolbox (mrVista). The high-resolution 3D-MDEFT anatomical data were used to manually segment the white-gray matter boundary in itkGray software, and 3D cortical surface and flat mesh models were created and realigned with the functional data by using mrVista. The functional images were corrected for motion in between and within scans^28^.

#### Population receptive field (pRF) mapping

For retinotopic mapping we used the population receptive field (pRF) method. The threshold of the explained variance (EV) was set to 15%, same with our human experiments and in agreement with previous studies^33,34^.

### MRI Stimuli

#### Stimulus presentation

Visual stimuli were displayed using a custom-made fiber-optic projection system at a resolution of 800 × 600 pixels (30 ° × 22.5 °) with a 60 Hz frame rate and mean luminance ∼50 cd/m^2^. Stimuli were centered on the approximate location of the fovea by using a modified fundus camera (Zeiss RC250). Animal eyes were fitted with appropriate contact lenses to ensure the stimulus remained in focus. At the beginning of each experiment a polar pattern with radius five degrees was presented monocularly to the left and right eyes in a block design (40 seconds ON – 40 seconds OFF) and the results were analysed online to select the eye with best alignment and the strongest responses. Further stimulus presentation was restricted to this eye.

#### Population RF (pRF) mapping

Moving square-checkerboard bars (100% contrast) with width 1° were moving in 0.5° steps every volume acquisition on the full screen extent sequentially in four different directions (Left-Right, Up-Down, Right-Left, Down-Up). Outside the bar aperture an isoluminant gray background was presented. Stimuli were generated using custom-made stimulus generation and presentation software (MRIstim). Each direction was presented twice for each scanning acquisition of 204 images. This was repeated 2-3 times.

#### Single unit RF (suRF) mapping

The suRF mapping binary checkerboard stimuli were generated with the same program creating the stimuli for the human subjects using Psychtoolbox-3^30^ in MATLAB (The Mathworks, Inc.). The only difference was that the block design ON-OFF periods were 24 and 36 seconds respectively and we used eight possible checker size conditions [0.25°, 0.5°, 1°, 2°, 4°, 6.5°, 13.0°, 22.5°]. Since the animals were under anesthesia no fixation spot was presented.

#### Electrophysiology data acquisition and preprocessing

Electrophysiological recordings were acquired from area V1 in two monkeys (M3, M4). Recording chambers were positioned stereotactically over the operculum with the aid of T1-weighted high-resolution 3D-MDEFT anatomical MRI images (0.5 mm isotropic). These images were used for targeting the region of interest, 3D-skull reconstruction and design of the implants to accurately fit the contours of the skull. A 5-axis CNC machine (Willemin-Macodel W428) was used to build these form-specific implants that resulted in an excellent fit between the implants and the underlying skull surface. In one animal (M4) the implant was constructed from medical-grade titanium, which precluded MRI measurements. In the second animal (M3), the chamber was constructed from polyether ether ketone plastic (TECAPEEK; Ensinger GmbH), which is MRI compatible and we could thus also acquire pRF and suRF measurements.

In M3 recordings were conducted primarily using custom made tetrodes (see previous work for detais^35^; N=42 experiments) and in some (N=4) a custom-made multi-channel linear probe with 10 platinum/iridium channels with inter-distance ∼150μm was used in parallel. After eliminating noisy channels with none or very little spiking activity and channels recorded from area V2 this resulted in N=35 datasets from area V1. In M4 recordings were conducted using either single-channel tungsten electrodes (FHC, Inc., Bowdoin, ME USA; N=8 experiments) or custom made multi-channel probes made with platinum/iridium wire (9 linearly arranged recording locations with inter-distances of ∼150μm, the 8th location was consisting of three nearby channels; N=10 experiments). After eliminating noisy channels with none or very little spiking activity and channels recorded from areas V2 and V4 this resulted in N=38 datasets that were classified as recordings from V1.

The electrodes were guided into the brain manually by custom designed adjustable microdrives. Electrophysiological activity was sampled at 32 kHz, digitized (16 bits), and stored using the Digital Lynx data acquisition system (Neuralynx). Multiunit activity was defined as the events that exceeded a predefined threshold (25 V) of the filtered, digitized signal (digital bandpass 600 Hz to 6 kHz). One animal (M4), was implanted with a scleral search coil^36,37^ and its eye movements were monitored on-line. In the second animal (M3), eye movements were monitored by an infrared eye-tracker (IView XTM Hi-Speed Primate, SMI). The behavioral aspects of the experiment were controlled using the QNX real-time operating system (QNX Software Systems Ltd).

#### Electrophysiology Stimuli

##### Stimulus Presentation

Visual stimuli were displayed using a dedicated graphics workstation (TDZ 2000; Intergraph Systems) running an OpenGL-based stimulation program (STIM). Stimuli were presented on a LCD monitor positioned at 1 m distance from the animals’ eyes (width: 60 cm, height: 34 cm) with a resolution of 1920 × 1080 and a refresh rate set to 100 Hz. The monitor was gamma corrected with a mean luminance of 22.2 cd/m2.

##### Reverse correlation RF mapping

To map the multiunit neuronal RFs we used the reverse correlation technique^21–25^. Stimuli were small square dots (black or white) presented on a gray background while the monkey fixated a small red spot with 0.2° diameter at the center of the monitor. The dots were positioned inside a rectangular grid with side dimensions of 0.5°-2.5° and location depending on a preliminary manual mapping of the RF location that was placed approximately in the center of the grid. Each dot was presented for 20 ms at pseudorandomized locations to have approximately equal number of presentations in each position inside the grid for each luminance (black and white).

##### Moving bar RF mapping

In one of the animals (M3), in parallel to the reverse correlation mapping, we also performed RF mapping with moving bars; this was similar to the pRF mapping performed in humans with the bars moving in eight different directions across the monitor in steps of 45°. The bar width was 0.5° and was moving by steps of 0.25° every 10 ms.

#### Electrophysiology Analysis

##### Eye-movement analysis

First, we calculated the time series of eye-velocities by differentiation of the position signals. Then, the horizontal and vertical angular velocities were independently thresholded at seven times their median-based SD to detect putative microsaccadic events. An event was classified as a microsaccade if the following additional criteria were satisfied: (1) it had a minimum duration of 8 ms, (2) it had an amplitude between 1 and 60 min of a degree, and (3) it had a maximum peak-velocity of 110° per second^38^. These parameters provided accurate detection of the microsaccades. In addition, the extracted microsaccades satisfied the main-sequence criterion and showed high correlation of amplitude and velocities. Fixation locations were extracted as the mean positions between saccades.

##### Reverse correlation RF mapping

The multiunit RFs at each recording location were reconstructed by iterative construction of spike histograms for each stimulus position. Each bin had a time span of 10 ms and in total we considered 12 bins (120 ms) post stimulus presentation. For each presentation of a dot (separately for black and white dots) the spikes following were binned according to the eye-movement corrected position of the dot. Eye-movement correction improved substantially the estimation of RFs in M4 that had an eye-coil. In contrast, in M3 (infrared eye-tracker) this procedure did not demonstrate improvements or even worsened RF reconstruction and thus it was not applied. Receptive field sizes with or without the correction in M3 were very similar with some changes in RF position that could reflect drifts of the infrared eye-tracker. From the visual field position histograms, we then created spatial maps for each time-bin. Separate maps for the dark and bright dots as well as the average of the two were created. These maps were converted to z-scores by subtracting the mean and dividing with the standard deviation of the first three time-bin maps (i.e. < 30 ms where no substantial response is expected or was observed). Then, the bin with maximal responses was identified and used for fitting a 2D-isometric Gaussian with its parameters reflecting RF position (x,y) and size (2σ). This procedure was performed for each map type (i.e. dark, bright, mean) and we selected the parameters of the model providing the best fit based on the coefficient of determination (R2). Examples of RF data and fits are presented in Fig. S10–S15.

##### Moving bar RF mapping

For the analysis of the moving bar RF mapping we have used a procedure similar to the pRF topographic mapping approach we proposed earlier for fMRI data^7^. For the analysis of the electrophysiological data an additional step was required before estimation of the RF. Specifically, a time delay reflecting the neuronal response latency was estimated so that the peaks of the responses when the bar was running in opposite directions were aligned. Also, in contrast to the fMRI analysis pipeline no hemodynamic deconvolution was necessary. All other steps of analysis were identical to the fMRI.

#### Modeling and Analysis

##### Single unit receptive field (suRF) modeling

In order to estimate the suRF size, the amplitudes of the responses (*β_BOLD_*) to the block design of checkerboard stimuli with different checker sizes *λ_chk_* presented in each block were estimated using a Generalized Linear Model (GLM). Only voxels that demonstrated GLM explained variance (EV) greater than 15% were further considered. The *β*’s for all conditions were then used to fit models of the general form of **Eq. 1** (see main manuscript) by using nonlinear least squares (*lsqcurvefit* function in Matlab; range of suRF-size between 0-10 degrees). These models use a static-nonlinearity to transform estimated neural responses (**Eq. 2** in the main manuscript) to BOLD signal. We considered several models (see Table 1).

For the purpose of modeling the receptive fields, we assume that neurons in primary visual cortex (V1) can be modeled as linear combinations of even (^*g_e_*^) and odd (^*g_o_*^) 2D-Gabor functions, described respectively with the following two equations:

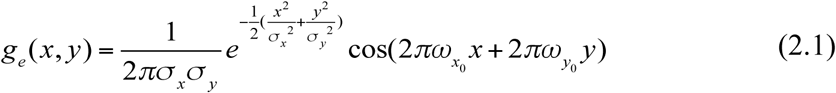

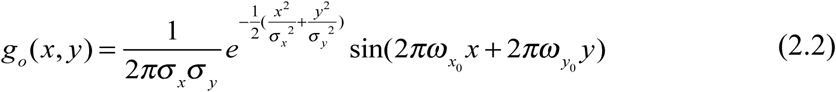

with *x* and*y* the Cartesian coordinates in the visual field, *σ_x_* and *σ_y_* defining the ellipsoid 2D Gaussian envelope and *ω*_*x*_0__, *ω*_*y*_0__ defining the center frequency and orientation. The spatial frequency sensitivities *ω_x_*,*ω_y_* of these Gabor filters can then be described by the respective Fourier transforms:

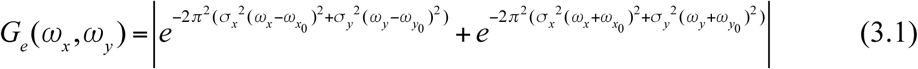

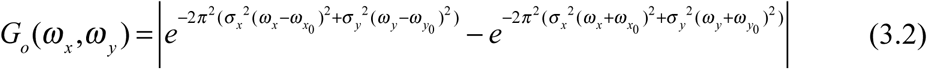

Further, we assume that the distribution of orientation selective cells within an fMRI voxel is approximately homogeneous, thus the spatial frequency sensitivity of voxels can be described relative to a single center frequency *ω*_0_ independent of orientation as follows:

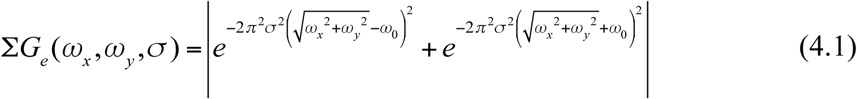

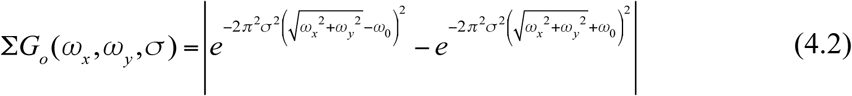

Thus, the spatial frequency sensitivity of the population of cells within a voxel can be estimated according to a linear combination of the equations 3.1 and 3.2. For simplicity, we take the average of the even and odd parts as follows: Σ*G*(*ω_x_*,*ω_y_*,*σ*) = (Σ*G_e_* + Σ*G_o_*)/2 (see also **Fig. S1**).

Then, for estimating the neural responses R (see Eq. 2 in the main manuscript), we integrated the product of ΣG with an analytical form of the stimulus frequency content S (see Fig. S2).

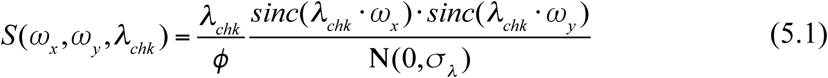

## Acknowledgments

This work was supported by the Max-Planck Society, the German Federal Ministry of Education and Research (BMBF; FKZ: 01GQ1002), the Plasticise Consortium (Project HEALTH-F2-2009-223524), and the NEI (R01 EY-024019 to SS). We thank Dr Yibin Shao for contributing in data collection and Joachim Werner for technical support.

## Author contributions

G.A.K and S.M.S conceived the idea. G.A.K, N.K.L, and S.M.S designed the experiments. G.A.K., A.P, and Q.L. collected the data and performed preliminary preprocessing and analysis. G.A.K. performed the computational modelling and final analysis. G.A.K wrote the manuscript All authors discussed the results and provided comments to the final manuscript.

## Supplementary Materials

**Table S1.** Parameter estimates [±95%CI] and statistics of linear regression on the relationship of receptive field sizes vs. eccentricity in human subjects as shown in Fig. 3 E-H. To test if intercept and slope are different for suRF and pRF we also performed analysis of covariance (ancova) on the population (last row H1-4). Second level comparisons (Tukey-Kramer) showed a significant difference of the intercepts (*P*=1.33×10^−2^) as well as the slopes (*P*=1.06×10^−10^).

| Subject | Model | Intercept | Slope | F-statistic | p-value |
| --- | --- | --- | --- | --- | --- |
| **H1** | **suRF** | $0.42 \pm 0.05$ | $0.06 \pm 0.01$ | 312.18 | $8.28 \times 10^{-52}$ |
| | **pRF** | $0.43 \pm 0.17$ | $0.24 \pm 0.02$ | 578.05 | $3.88 \times 10^{-79}$ |
| **H2** | **suRF** | $0.60 \pm 0.12$ | $0.05 \pm 0.01$ | 44.85 | $1.02 \times 10^{-10}$ |
| | **pRF** | $1.22 \pm 0.41$ | $0.45 \pm 0.05$ | 330.96 | $1.22 \times 10^{-50}$ |
| **H3** | **suRF** | $0.81 \pm 0.24$ | $0.15 \pm 0.04$ | 46.82 | $1.03 \times 10^{-10}$ |
| | **pRF** | $1.01 \pm 0.49$ | $0.44 \pm 0.09$ | 91.25 | $6.37 \times 10^{-18}$ |
| **H4** | **suRF** | $0.63 \pm 0.13$ | $0.07 \pm 0.02$ | 60.28 | $2.49 \times 10^{-13}$ |
| | **pRF** | $0.72 \pm 0.36$ | $0.38 \pm 0.05$ | 255.00 | $2.05 \times 10^{-39}$ |
| **H1-4** | **suRF** | $0.69 \pm 0.07$ | $0.05 \pm 0.01$ | 139.47 | $1.96 \times 10^{-30}$ |
| | **pRF** | $0.94 \pm 0.19$ | $0.33 \pm 0.02$ | 727.71 | $3.73 \times 10^{-124}$ |

**Table S2.** Parameter estimates [±95%CI] and statistics of linear regression on the relationship of receptive field sizes vs. eccentricity in monkey subjects as shown in Fig. 4 E-G. To test if intercept and slope are different for suRF and pRF we also performed analysis of covariance (ancova) on the population (last row M1-2). Second level comparisons (Tukey-Kramer) between the pRF and suRF showed a significant difference of the intercepts (*P*=9.56×10^−10^) as well as the slopes (*P*=9.56×10^−10^). In contrast, the difference of the suRF with electrophysiology was not-significant for the intercepts (*P*=0.47) or the slopes (*P*=0.98).

| Subject | Model | Intercept | Slope | F-statistic | p-value |
| --- | --- | --- | --- | --- | --- |
| **M1** | **suRF** | $0.69 \pm 0.17$ | $0.03 \pm 0.02$ | 13.66 | $2.52 \times 10^{-4}$ |
| | **pRF** | $1.97 \pm 0.46$ | $0.13 \pm 0.05$ | 25.13 | $8.27 \times 10^{-7}$ |
| **M2** | **suRF** | $0.39 \pm 0.04$ | $0.06 \pm 0.01$ | 365.00 | $4.05 \times 10^{-63}$ |
| | **pRF** | $1.29 \pm 0.16$ | $0.20 \pm 0.03$ | 241.35 | $1.44 \times 10^{-45}$ |
| **M1-2** | **suRF** | $0.42 \pm 0.05$ | $0.06 \pm 0.01$ | 325.61 | $9.37 \times 10^{-63}$ |
| | **pRF** | $1.39 \pm 0.15$ | $0.19 \pm 0.02$ | 319.08 | $1.07 \times 10^{-61}$ |
| **M3-4** | **ephys** | $0.16 \pm 0.05$ | $0.08 \pm 0.02$ | 63.36 | $1.99 \times 10^{-11}$ |

**Fig. S1.**
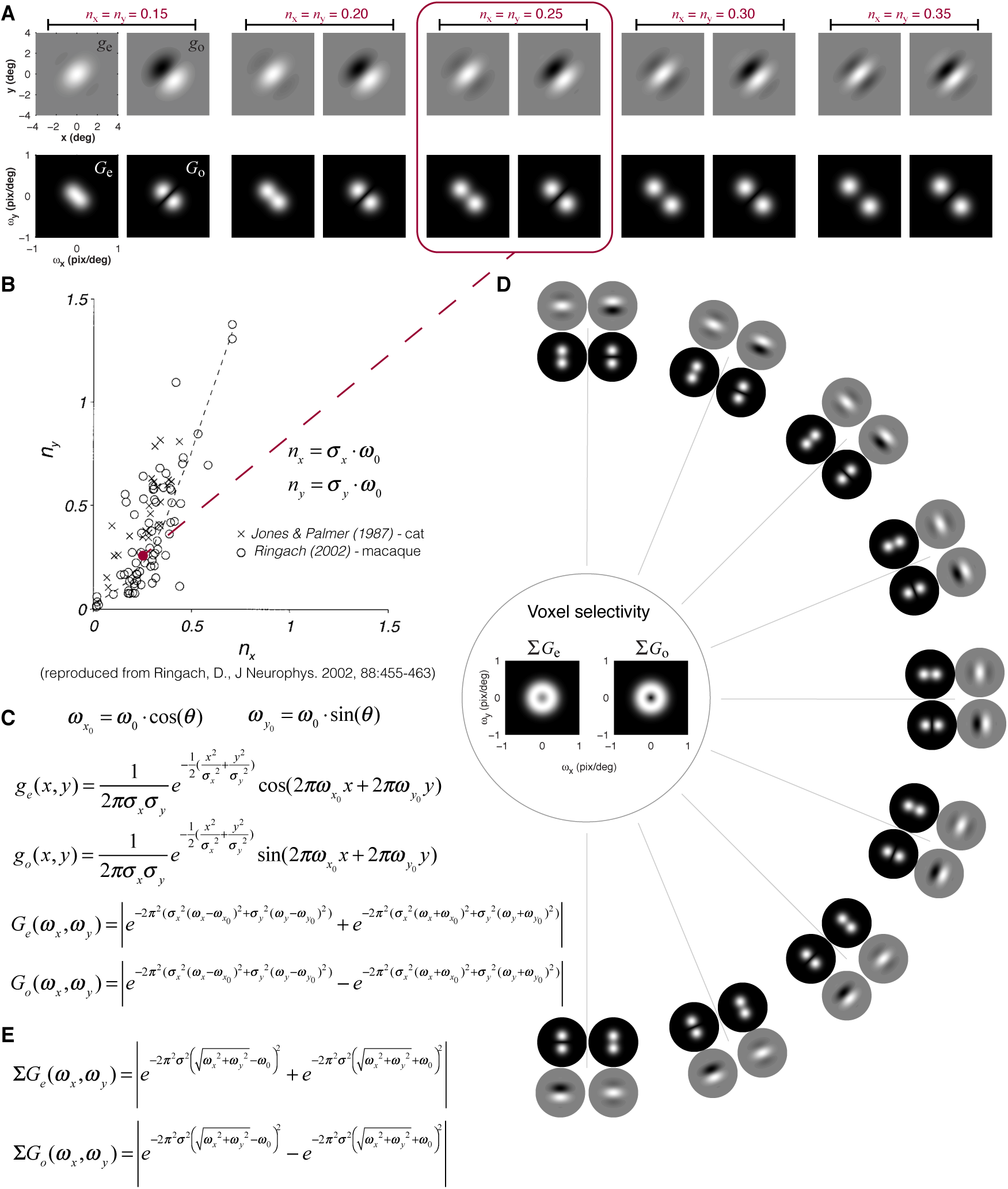
Voxel spatial frequency selectivity based on Gabor-like receptive fields^36,37^. (A) Examples of even (*g*e) and odd (*g*o) symmetric 2D-Gabor pairs in Cartesian x-y space (top) and their respective spatial frequency selectivity in frequency space (bottom) for different values of parameter *n_x_* that controls the shape of the Gabor (number of subunits) by the relationship between the frequency of the sinusoid and the standard deviation of the Gaussian envelope. (B) We chose to use *n_x_* = *n_y_* = 0.25 that represents a central tendency of data recorded from cats^38^ and monkeys^24^. (C) Equations of the Gabors (*g_e_*, *g_o_*) and their spatial frequency selectivity (*G_e_*, *G_0_*) as shown in A. (D) Assuming a homogeneous representation of Gabor orientations within a voxel we can estimate the spatial frequency selectivity of the voxel independent of orientation (S*G_e_*, S*G_o_*). (E) Equations that estimate the spatial frequency selectivity of the voxel depending on the envelope standard deviation *σ* (note that ω_0_ is also depending on *σ* based on *n_x_*).

**Fig. S2.**
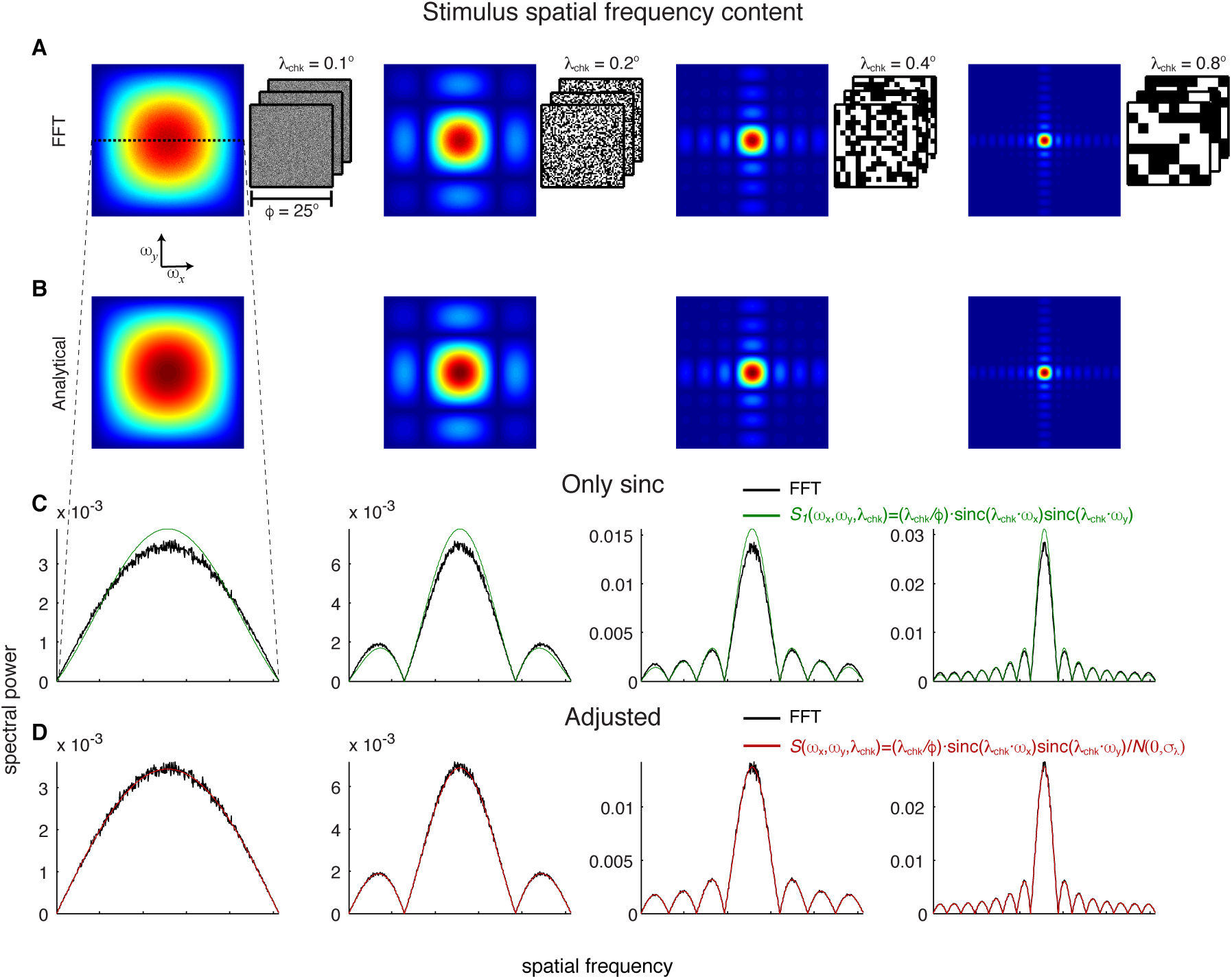
Stimulus and its spatial frequency content. Stimuli were full-field binary checkerboard patterns switching randomly with a frame rate of 30 Hz in blocks of 10 seconds. We manipulated the average spatial frequency content by changing the size of the square checkers with edge *λ*_chk_. In (A) a cartoon of three stimulus frames is shown next to the average spatial frequency content estimated with the fast Fourier-transform (FFT) for four example conditions [*λ*_chk_ = 0.1°, 0.2 °, 0.4 °, 0.8 °]. Panel (B) shows the analytical estimation of the same spatial frequencies using an adjusted 2D sinc function. Since the 2D sinc function corresponds to the transformation of a single checker, we adjusted it by dividing with an additional Gaussian of standard deviation σ_1_ (depending on the smallest possible checker) to take into account the full checkerboard. (C-D) demonstrate the difference between the 2D-sinc and adjusted 2D-sinc estimators along the horizontal axis demonstrating that the adjusted version nicely estimates the correct frequency content.

**Fig. S3.**
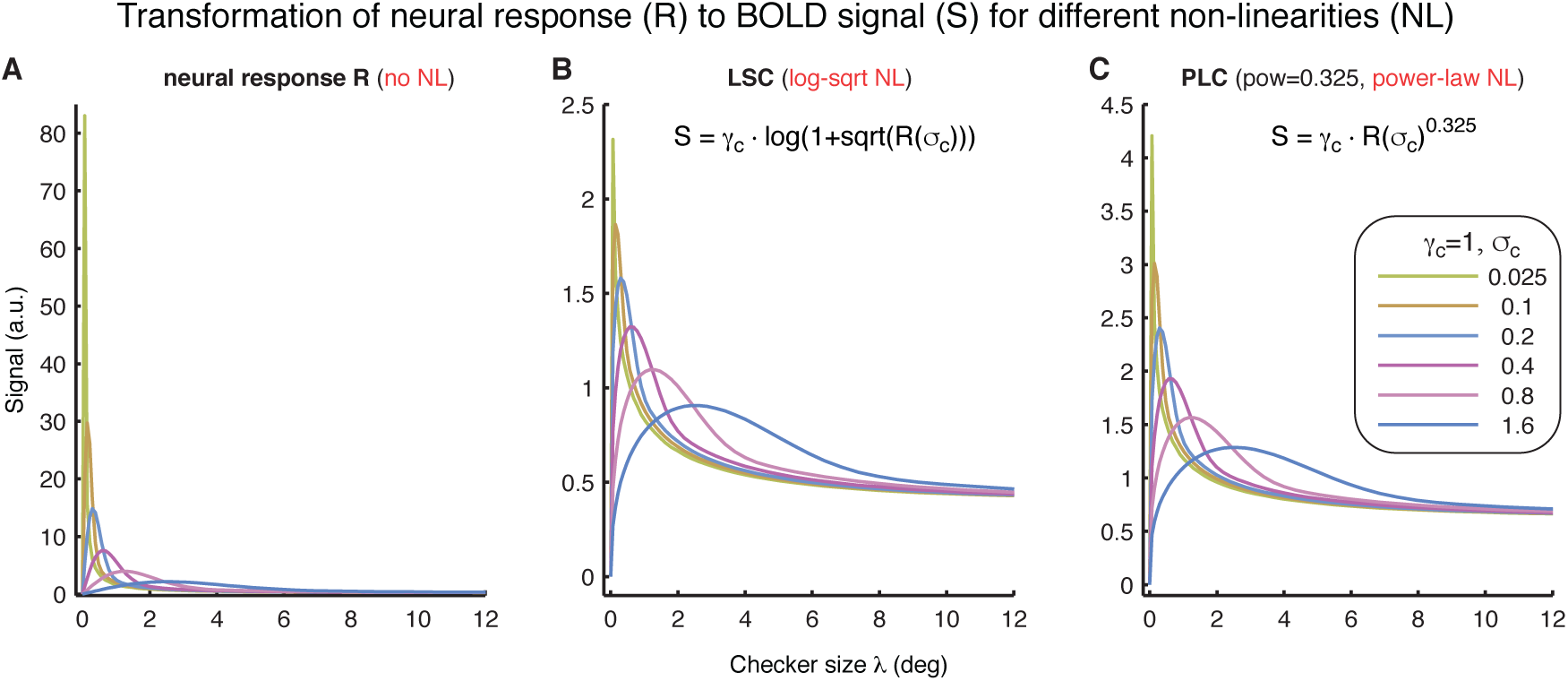
Transformation of neural responses to BOLD signal using different static non-linearities. In (A) the raw neural response (R) is shown. In (B) the LSC model using a log-square root non-linearity is presented for different receptive field sizes σ_c_ (see legend). (C) similar to B but for the PLC model which incorporates a power-law nonlinearity.

**Fig. S4.**
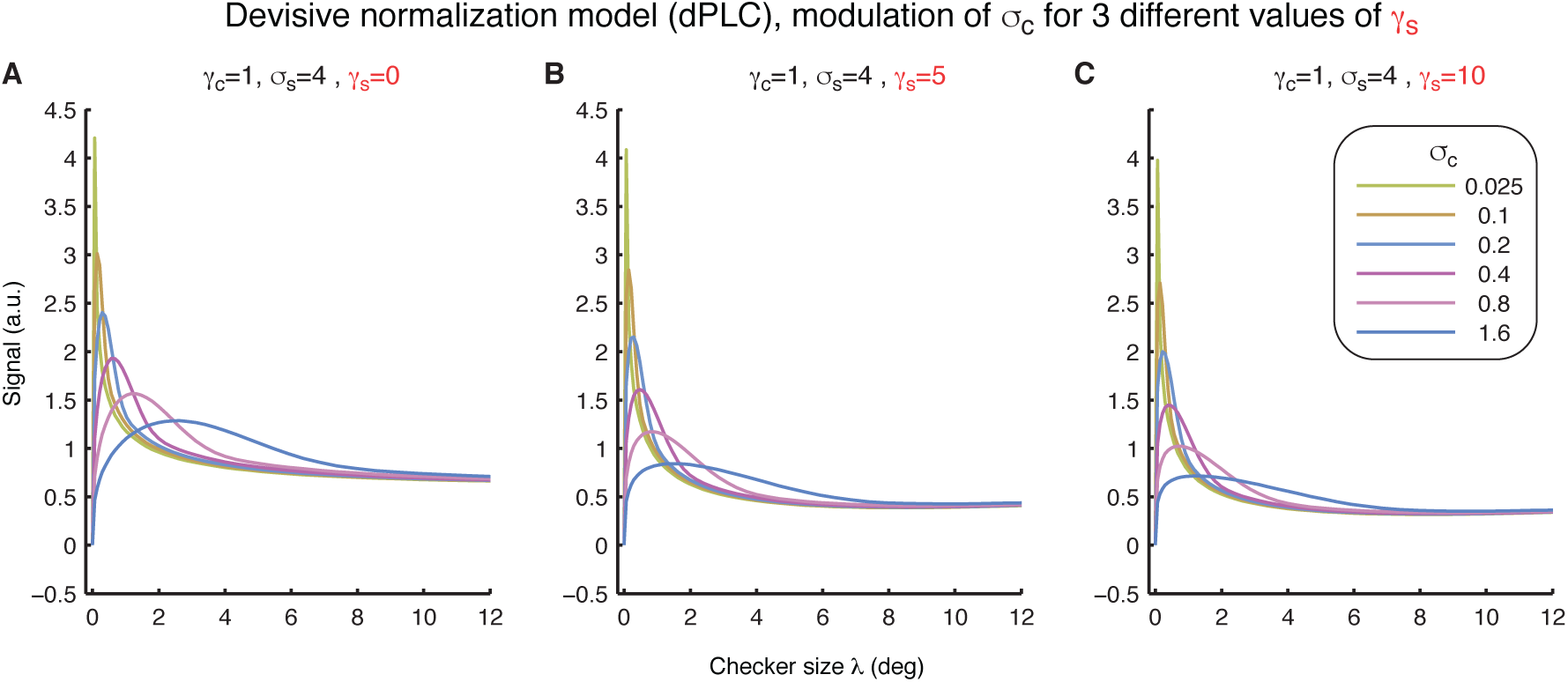
Demonstration of the effects of the amplitude of divisive normalization (dPLC model with σ_s_=4°) as given by changing the gain parameter *γ*_s_.

**Fig. S5.**
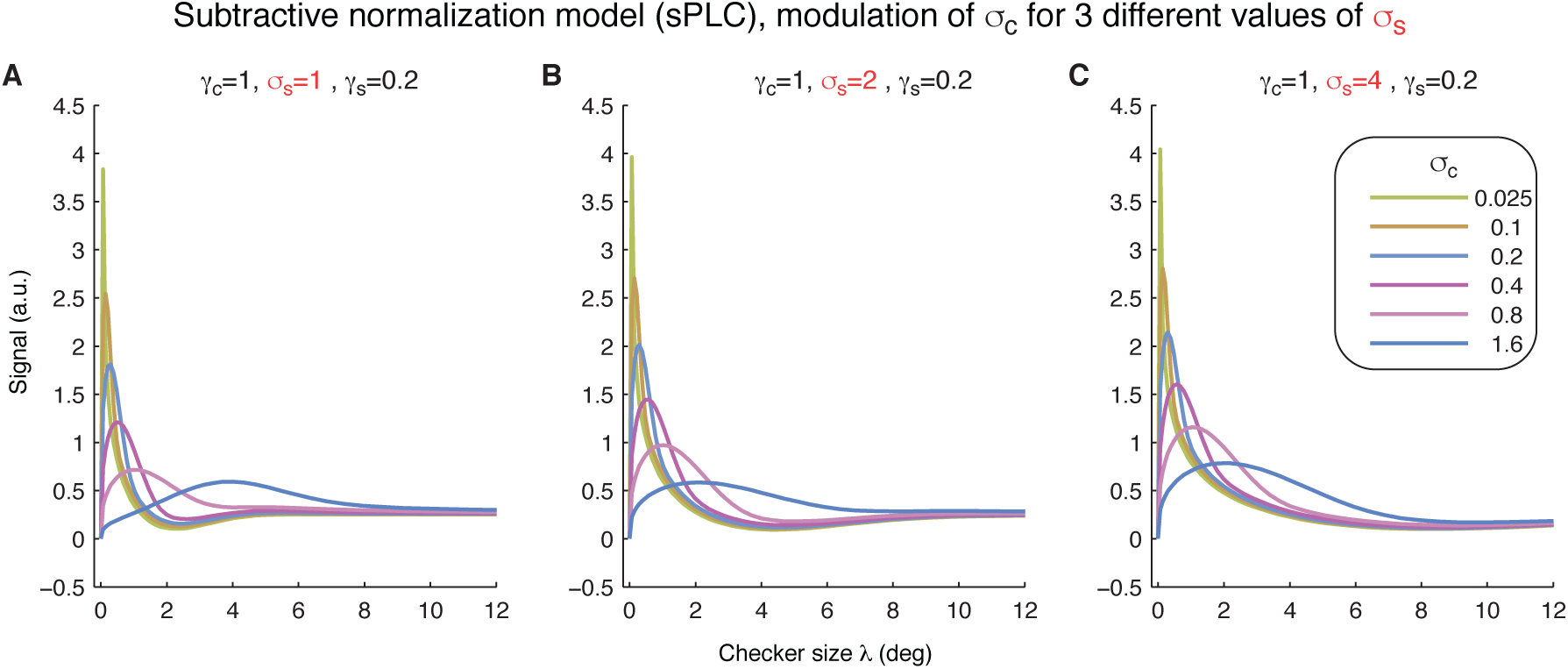
Demonstration of the effects of receptive field size of subtractive normalization (sPLC model with *γ*_s_=0.2) as given by changing σ_s_.

**Fig. S6.**
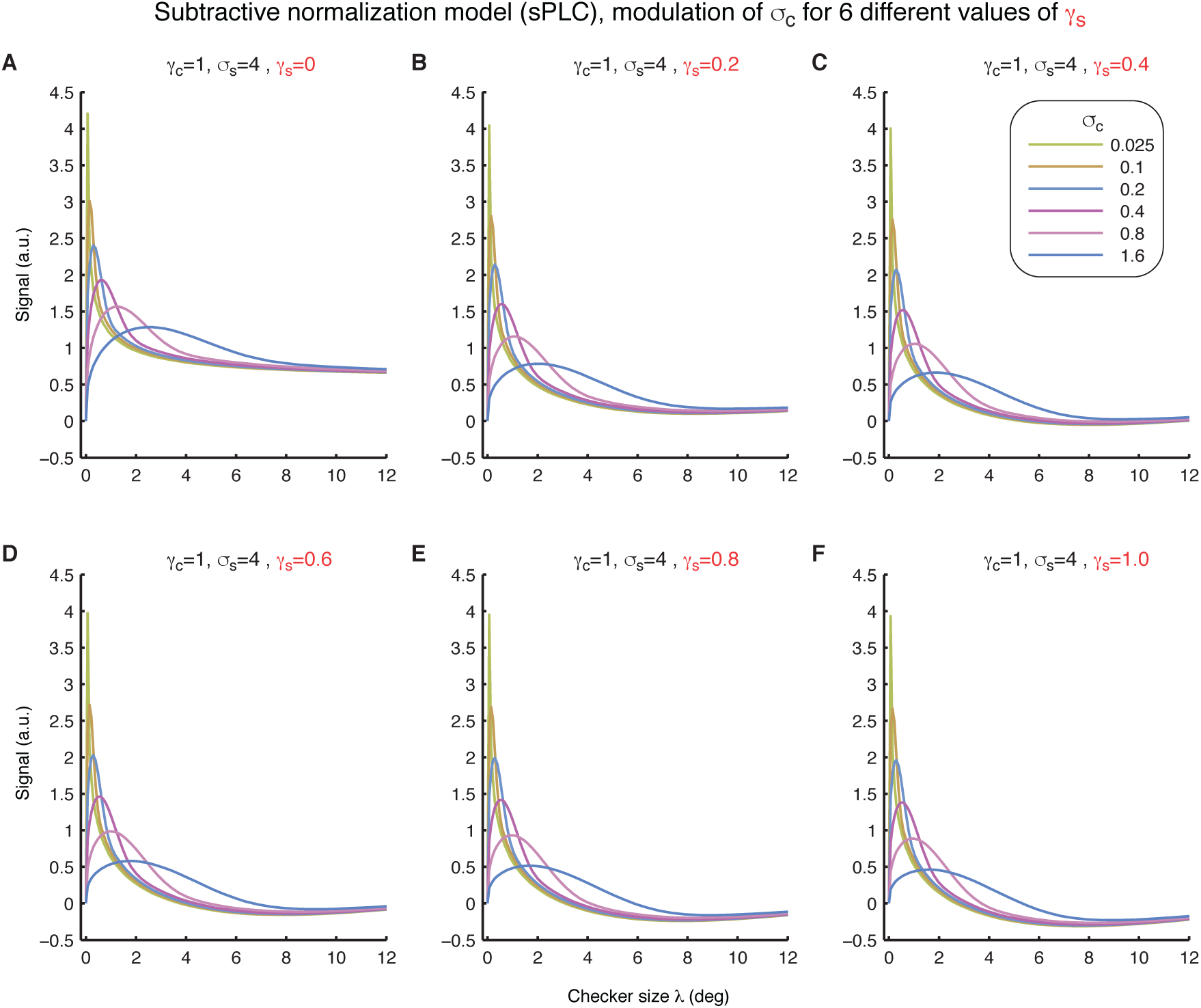
Demonstration of the effects of the amplitude of subtractive normalization (sPLC model with σ_s_=4°) as given by changing the gain parameter *γ*_s_.

**Fig. S7.**
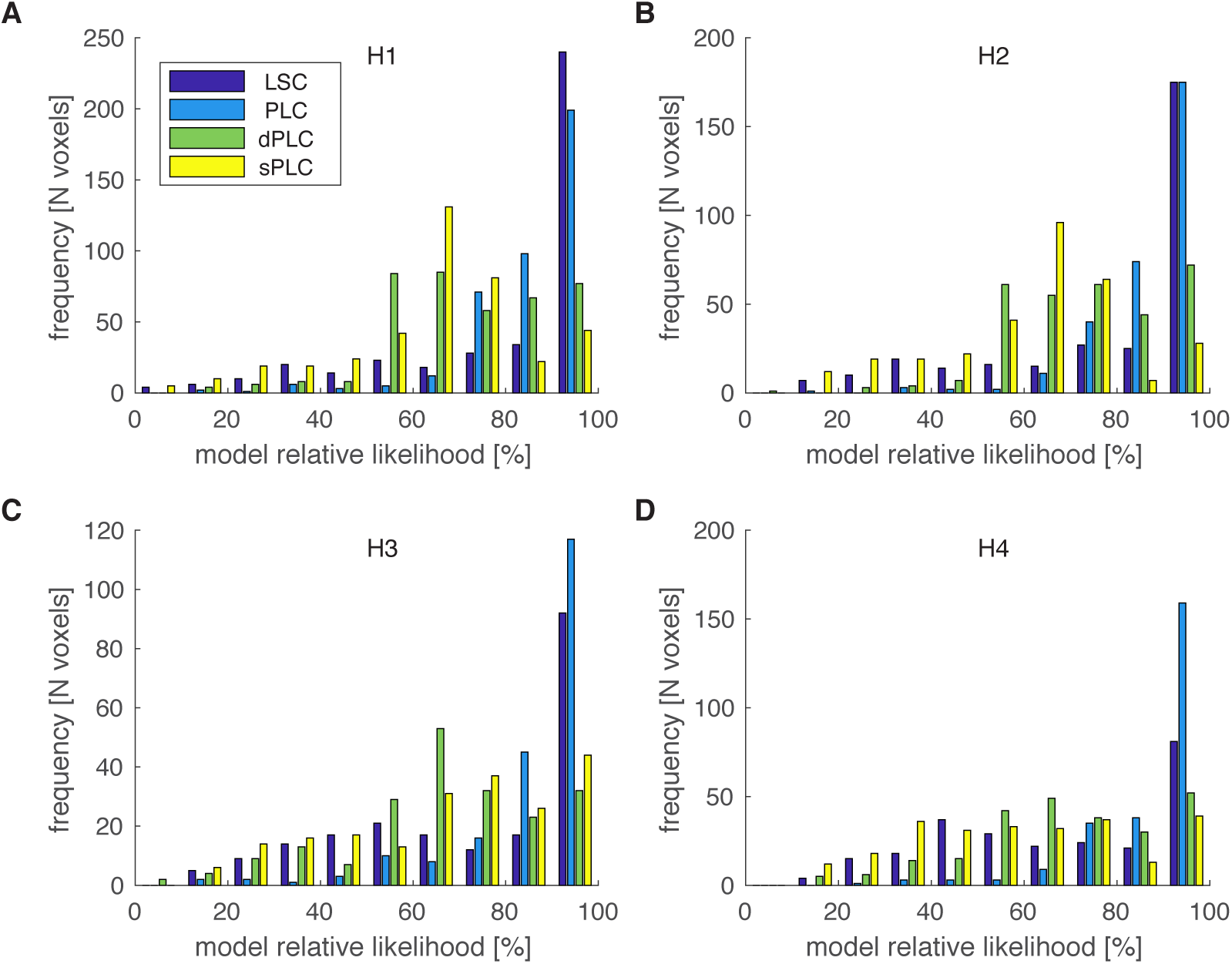
Model relative likelihood distributions as derived from AIC scores ^17^ across voxels of four different models: LSC, PLC, dPLC, and sPLC (see Table S1 for details on the models) Each panel A-D presents the results of each human subject H1-H4 respectively.

**Fig. S8.**
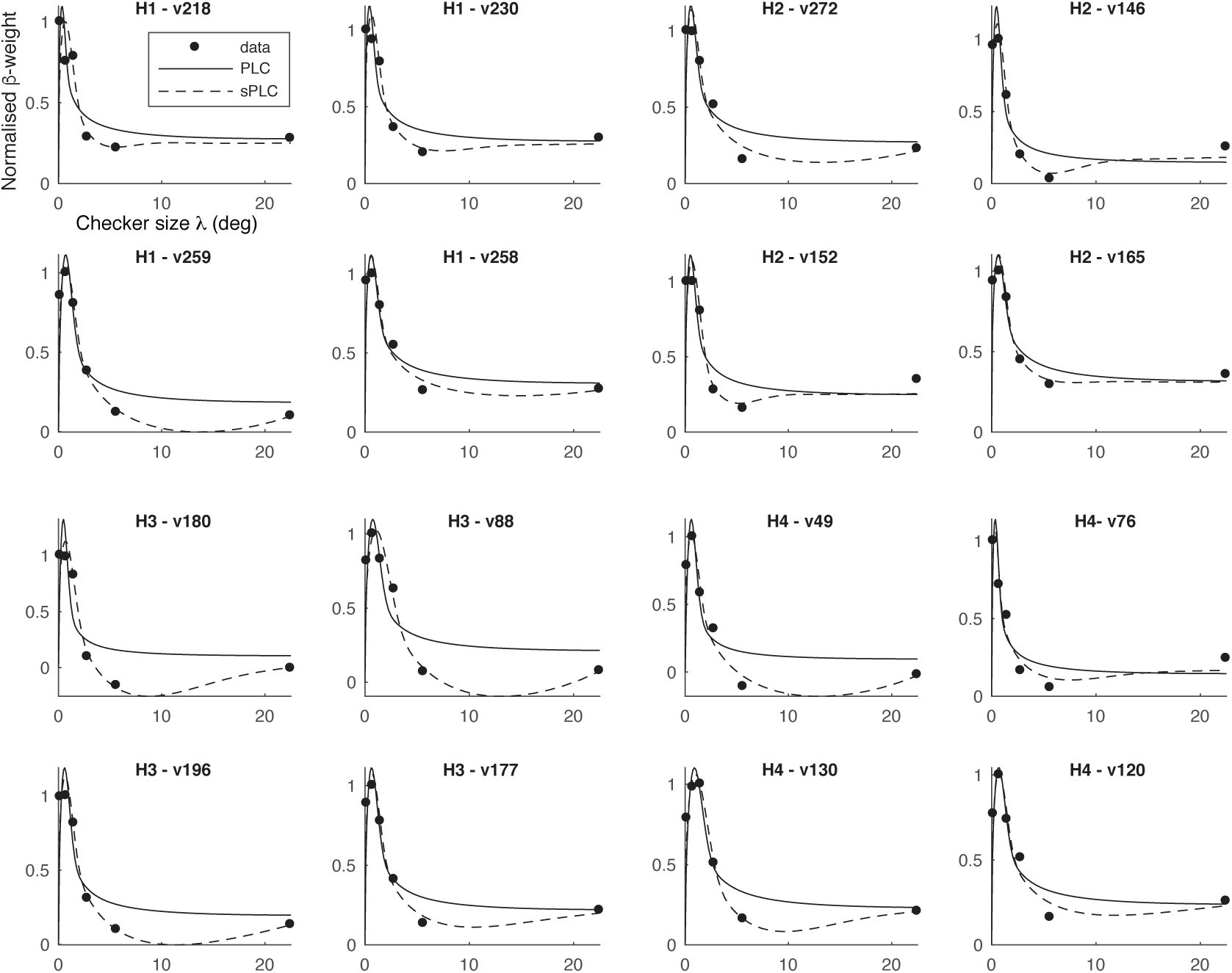
Selected example voxels for which the subtractive normalization model (sPLC) outperforms the no-normalization model (PLC) based on Akaike’ s information criterion (AIC). Note that the responses as reflected in the normalized β-weights demonstrate suppression for intermediate checker sizes and that this could not be fit by the PLC model (see Fig. S3 C for PLC model behavior). H1-H4 reflect the different subjects.

**Fig. S9.**
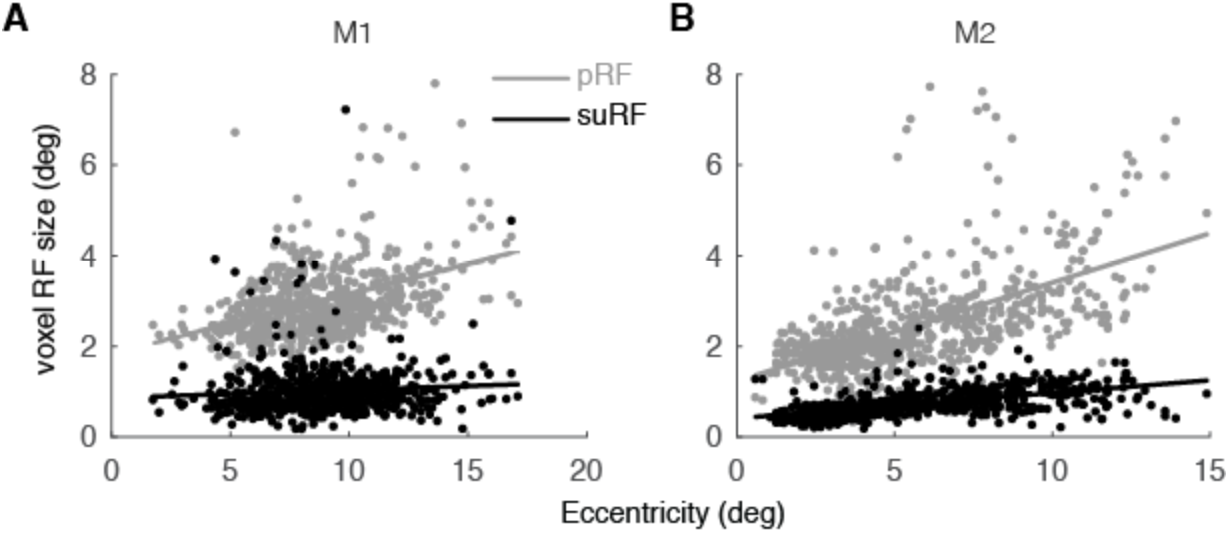
FMRI estimated RF sizes versus eccentricity for explained variance (EV) threshold of 10% instead of 15% used throughout the manuscript (see Fig. 4 for the respective plots in the monkeys). (A-B) pRF (gray) and suRF (black) size as a function of eccentricity for monkeys M1-M2 respectively. Each dot represents a voxel. Lines represent linear regression fits. Note that data points extent more towards the fovea in comparison to Fig. 4 but the general relationships remain the same. We conjecture that the limited significance of activation in the fovea in monkeys in contrast to humans could be explained: a) by the known difficulty to activate foveal regions in anesthetized macaques, b) increased effects of surround suppression observed in the fovea during anesthesia and no eye jittering. Importantly, a large proportion of foveal voxels in monkey demonstrated suppression (see Fig. 4A) and thus could be better fitted with the subtractive normalization model (PLC) like in Fig. S8.

**Fig. S10.**
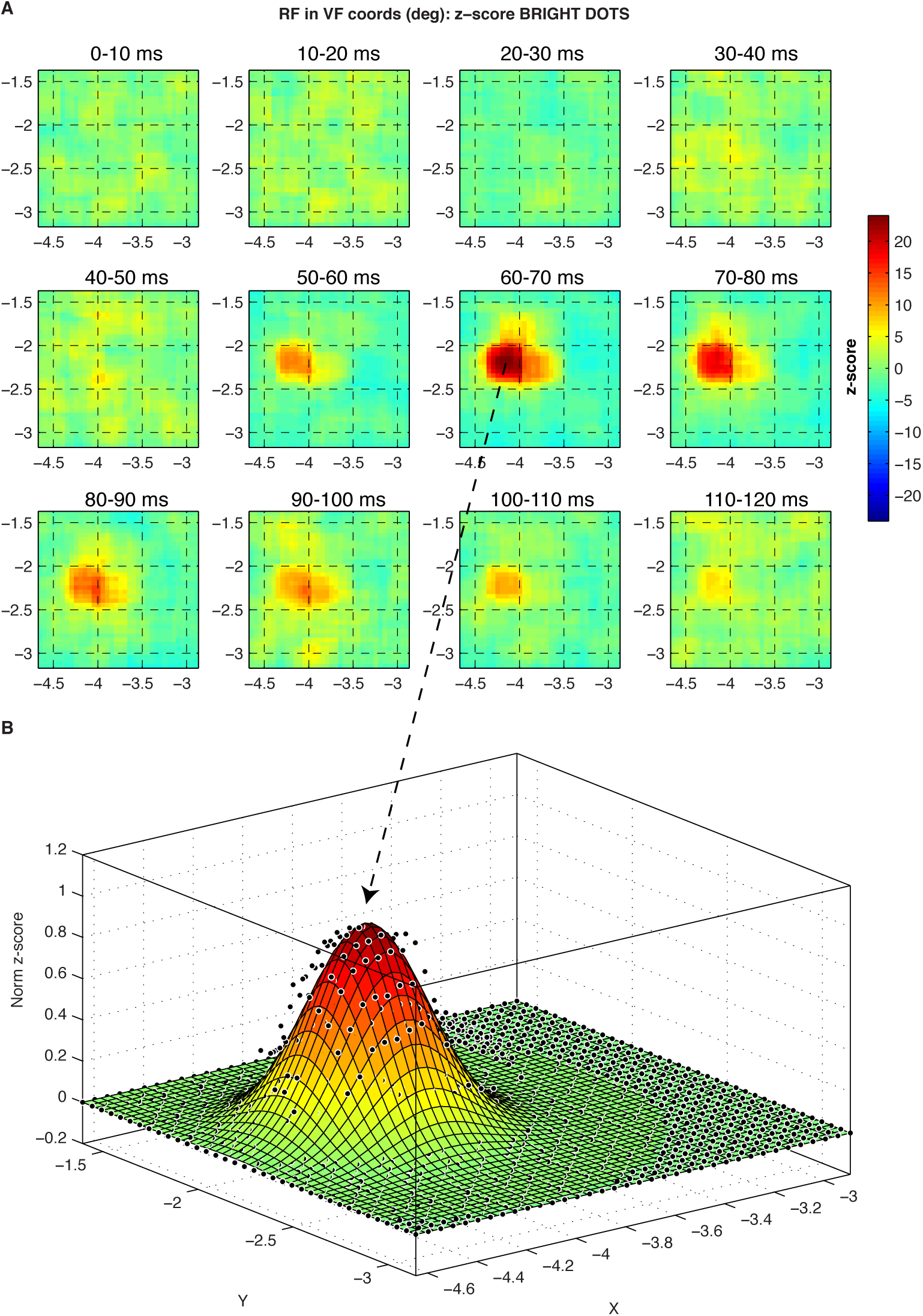
Example of reverse correlation receptive field recorded by electrophysiological recordings in V1. In panel A, the spatiotemporal RF of multi-unit activity is presented using two-dimensional segments of the visual fields that reflect time windows of 10 ms. Time zero represents the recording of a spike and positive times run backward so that in each panel the average stimulus during a window of time before the spike is presented. In panel B, we show the fit of a 2D isometric Gaussian to the data. The parameter σ of the Gaussian is assumed to be proportional to RF size (2σ).

**Fig. S11.**
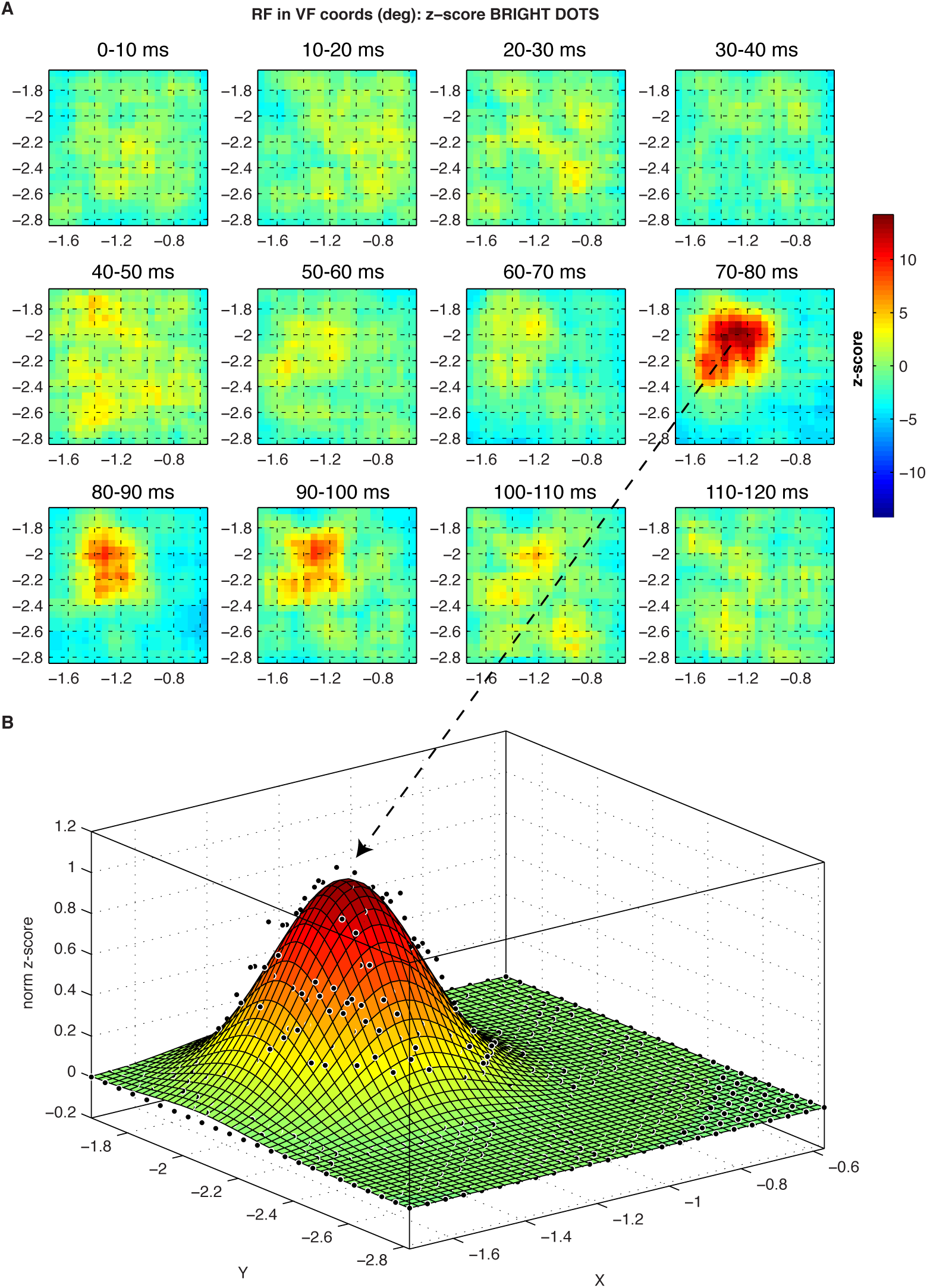
Example of reverse correlation receptive field recorded by electrophysiological recordings in V1. In panel A, the spatiotemporal RF of multi-unit activity is presented using two-dimensional segments of the visual fields that reflect time windows of 10 ms. Time zero represents the recording of a spike and positive times run backward so that in each panel the average stimulus during a window of time before the spike is presented. In panel B, we show the fit of a 2D isometric Gaussian to the data. The parameter σ of the Gaussian is assumed to be proportional to RF size (2σ).

**Fig. S12.**
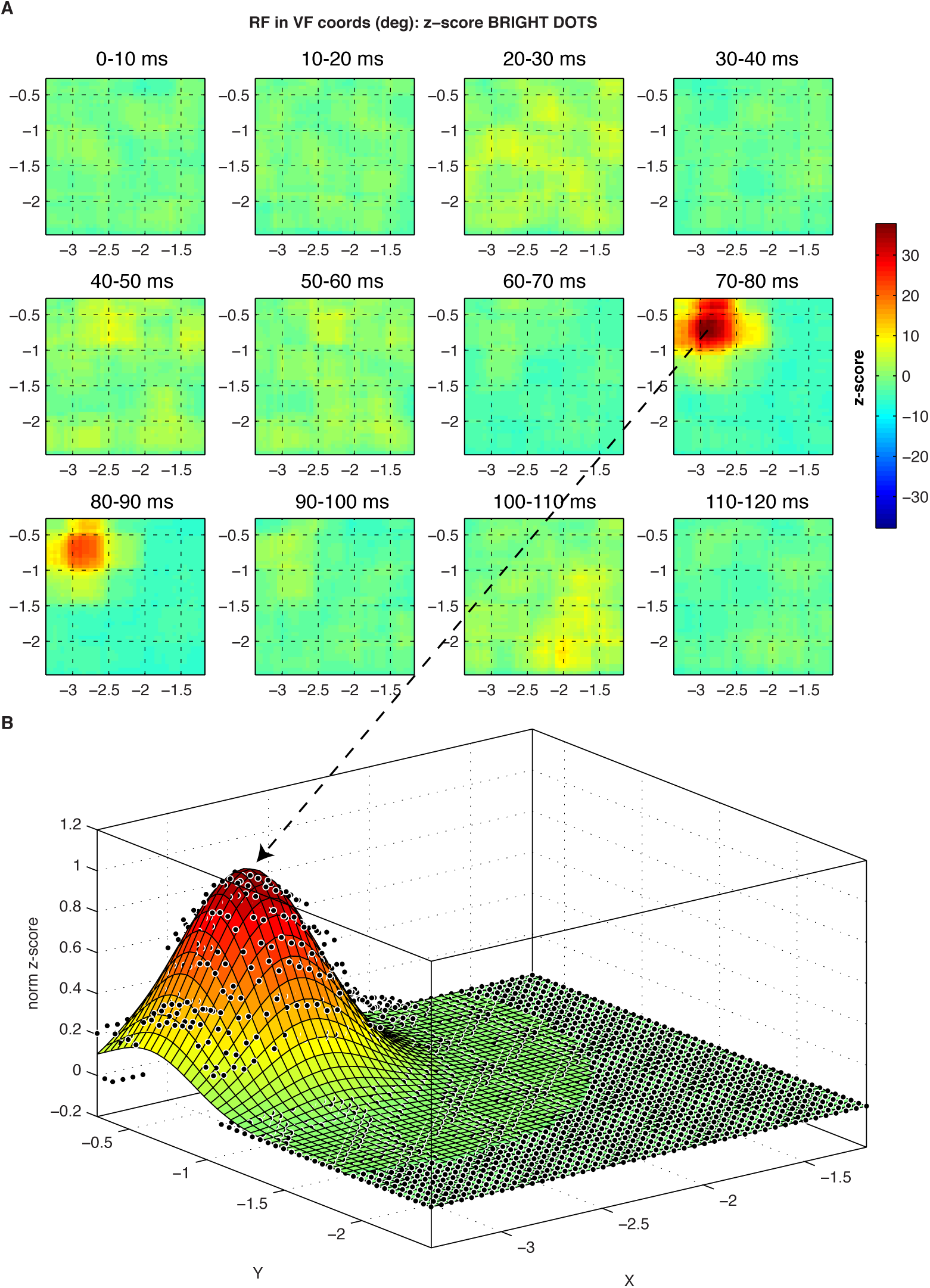
Example of reverse correlation receptive field recorded by electrophysiological recordings in V1. In panel A, the spatiotemporal RF of multi-unit activity is presented using two-dimensional segments of the visual fields that reflect time windows of 10 ms. Time zero represents the recording of a spike and positive times run backward so that in each panel the average stimulus during a window of time before the spike is presented. In panel B, we show the fit of a 2D isometric Gaussian to the data. The parameter σ of the Gaussian is assumed to be proportional to RF size (2σ).

**Fig. S13.**
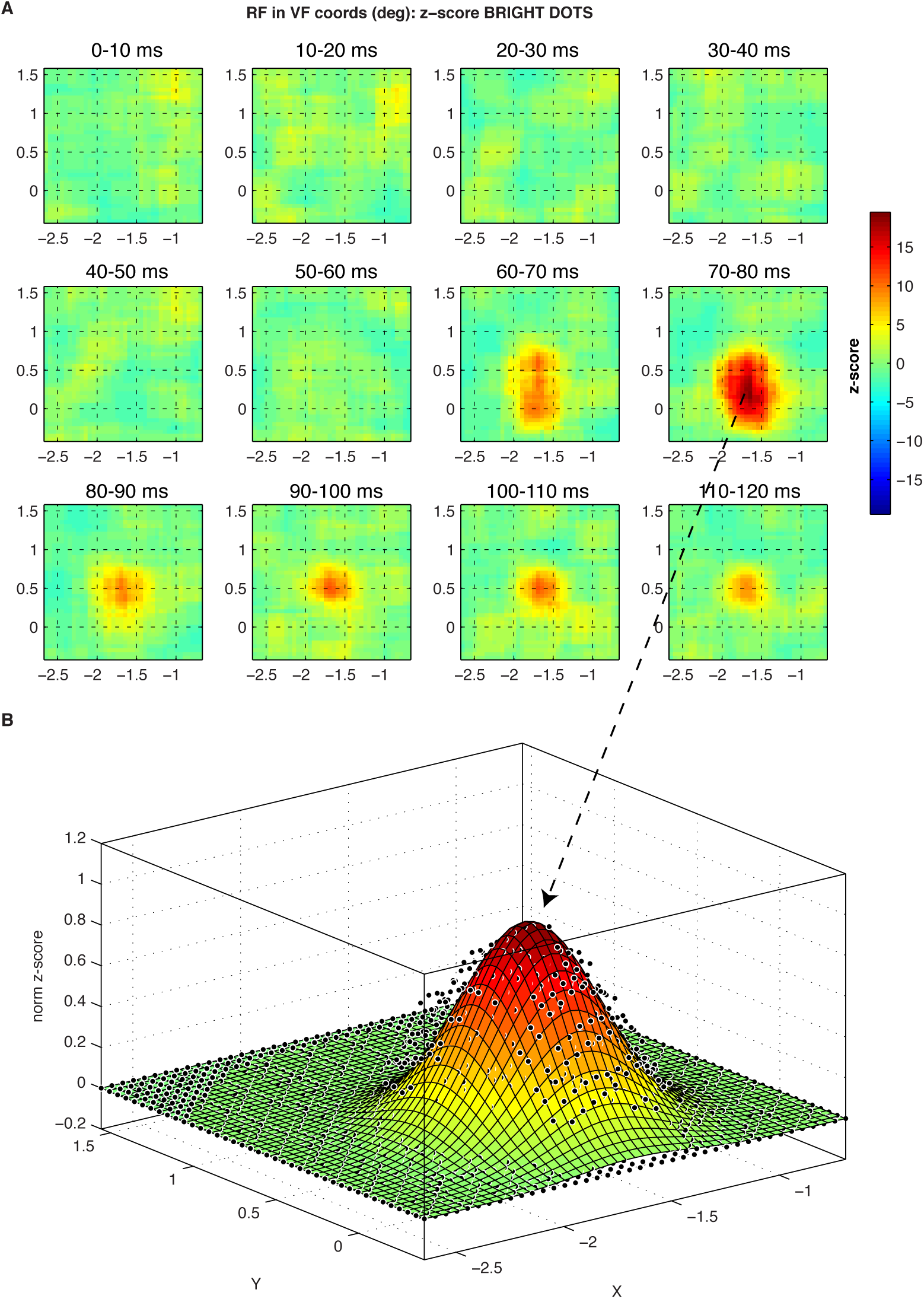
Example of reverse correlation receptive field recorded by electrophysiological recordings in V1. In panel A, the spatiotemporal RF of multi-unit activity is presented using two-dimensional segments of the visual fields that reflect time windows of 10 ms. Time zero represents the recording of a spike and positive times run backward so that in each panel the average stimulus during a window of time before the spike is presented. In panel B, we show the fit of a 2D isometric Gaussian to the data. The parameter σ of the Gaussian is assumed to be proportional to RF size (2σ).

**Fig. S14.**
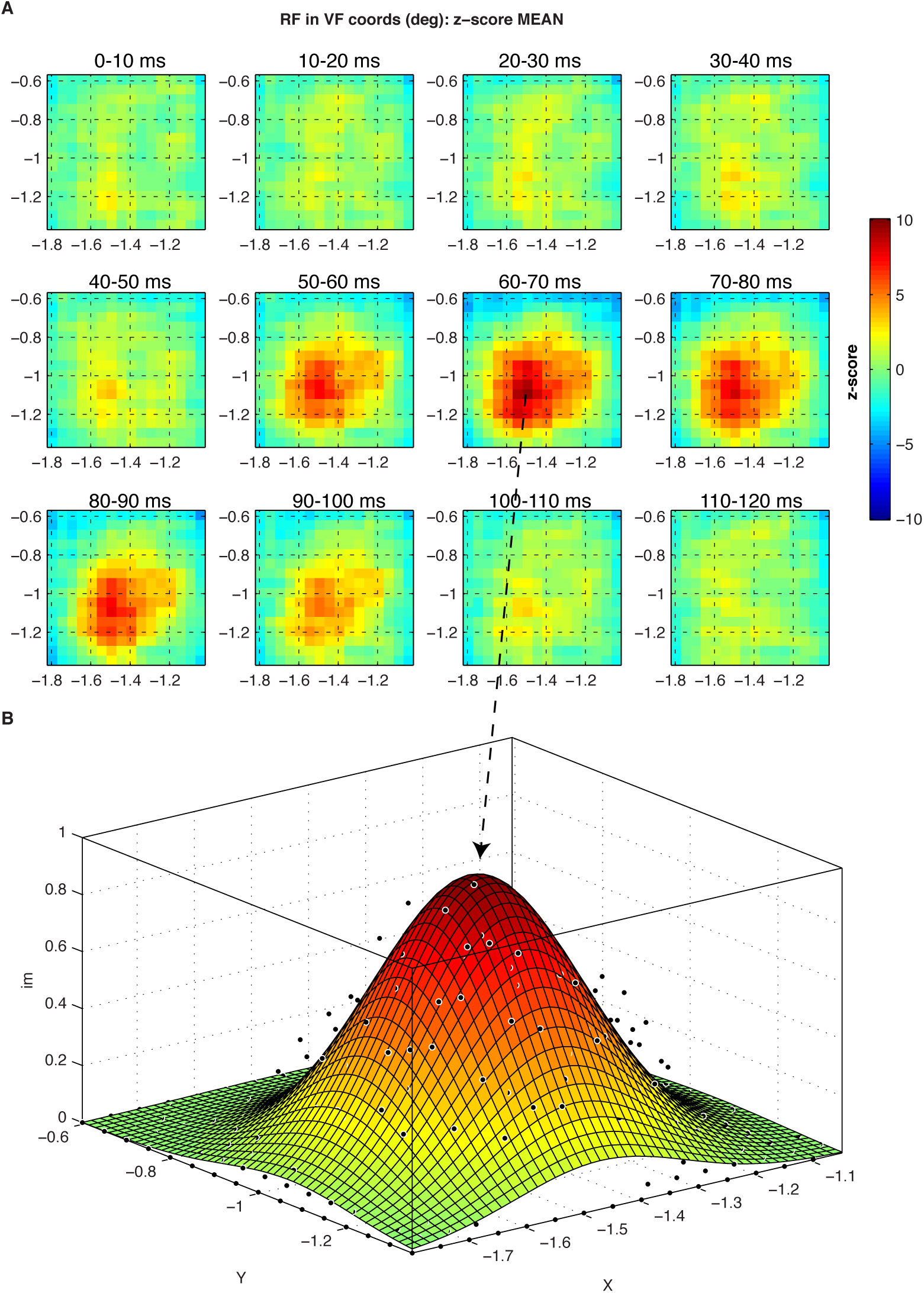
Example of reverse correlation receptive field recorded by electrophysiological recordings in V1. In panel A, the spatiotemporal RF of multi-unit activity is presented using two-dimensional segments of the visual fields that reflect time windows of 10 ms. Time zero represents the recording of a spike and positive times run backward so that in each panel the average stimulus during a window of time before the spike is presented. In panel B, we show the fit of a 2D isometric Gaussian to the data. The parameter σ of the Gaussian is assumed to be proportional to RF size (2σ).

**Fig. S15.**
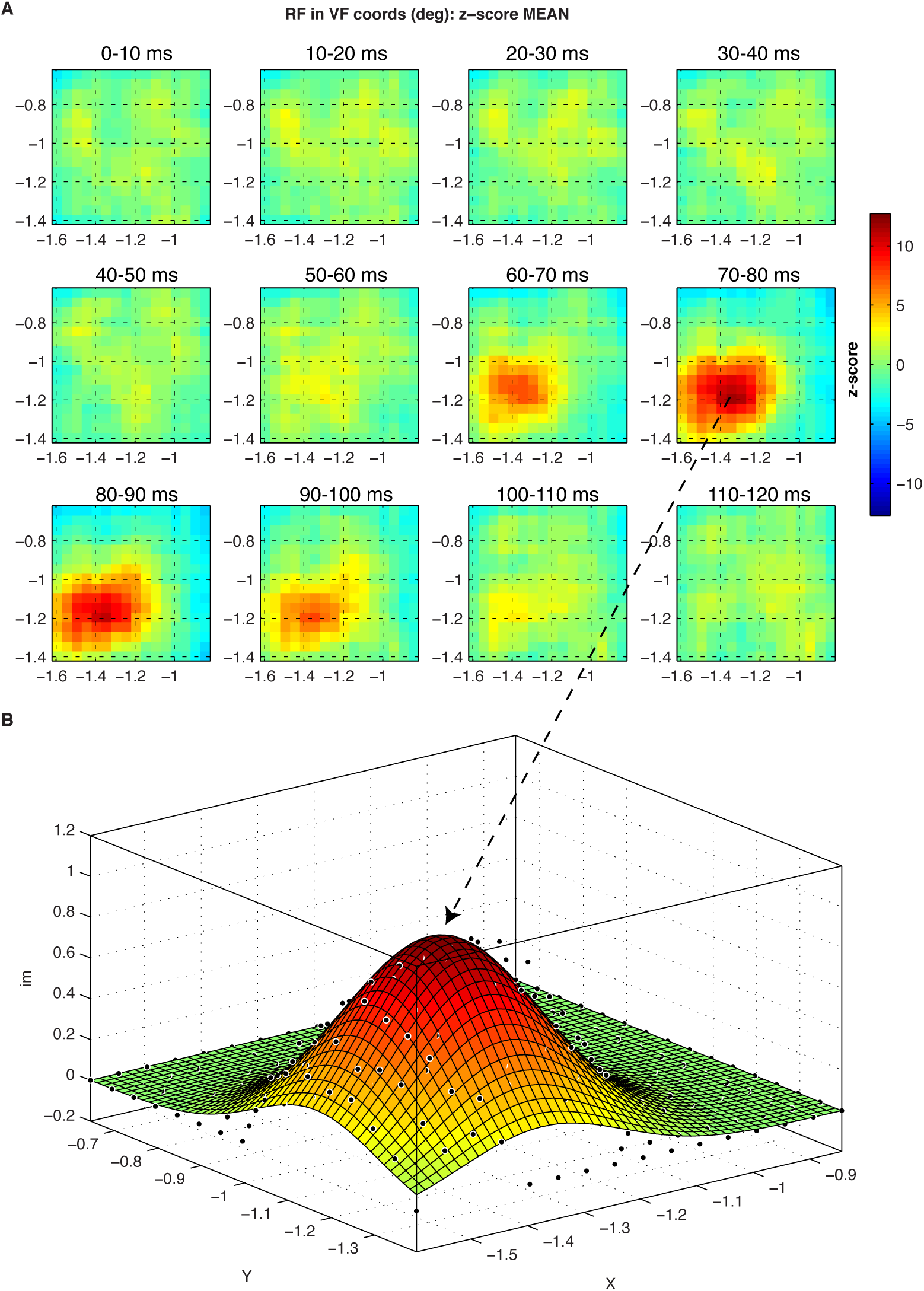
Example of reverse correlation receptive field recorded by electrophysiological recordings in V1. In panel A, the spatiotemporal RF of multi-unit activity is presented using two-dimensional segments of the visual fields that reflect time windows of 10 ms. Time zero represents the recording of a spike and positive times run backward so that in each panel the average stimulus during a window of time before the spike is presented. In panel B, we show the fit of a 2D isometric Gaussian to the data. The parameter σ of the Gaussian is assumed to be proportional to RF size (2σ).

